# Exploring associations between the teat apex metagenome and *Staphylococcus aureus* intramammary infection risk in primiparous cows under organic directives

**DOI:** 10.1101/2023.09.12.557435

**Authors:** C. J. Dean, F. Peña-Mosca, T. Ray, T. J. Wehri, K. Sharpe, A.M. Antunes, E. Doster, L. Fernandes, V. F. Calles, C. Bauman, S. Godden, B. Heins, P. Pinedo, V. S. Machado, L. S. Caixeta, N. R. Noyes

## Abstract

The primary objective of this study was to identify associations between teat apex microbiome and *Staphylococcus aureus* intramammary infection (IMI) risk in primiparous cows during the first 5 weeks after calving. We performed a case-control study using shotgun metagenomics of the teat apex and culture-based milk data collected longitudinally from 710 primiparous cows on 5 organic dairy farms. We observed a strong association between *S. aureus* DNA in the metagenomic teat apex data prior to parturition and the odds of *S. aureus* IMI after parturition (OR = 38.9, 95% CI: 14.84-102.21). Differential abundance analysis confirmed this association, with cases having a 23.8 higher log fold change (LFC) in abundance of *S. aureus* in their samples compared to controls. Of the most prevalent microorganisms in controls, those associated with a lower risk of post-calving *S. aureus* IMI included *Microbacterium* phage Min 1 (OR = 0.37, 95% CI: 0.25-0.53), *Corynebacterium efficiens* (OR = 0.53, 95% CI: 0.30-0.94), *Kocuria polaris* (OR = 0.54, 95% CI: 0.35-0.82), *Micrococcus terreus* (OR = 0.64, 95% CI: 0.44-0.93) and *Dietzia alimentaria* (OR = 0.45, 95% CI: 0.26-0.75). Microcin B17 was the most prevalent antibacterial peptide on the teat apex of cases and controls (99.7% in both groups). The predicted abundance of Microcin B17 was also higher in cases compared to controls (LFC 0.26). Cow and farm random effects often explained a large proportion of the observed variability in the teat apex microbiome, suggesting that our results need to be interpreted within the context of the random effects.

**IMPORTANCE:** Intramammary infections (IMI) caused by *Staphylococcus aureus* remain an important problem for the organic dairy industry. The microbiome on the external skin of the teat apex may play a role in mitigating *S. aureus* IMI risk, in particular the production of antimicrobial peptides (AMPs) by commensal microbes. However, current studies of the teat apex microbiome utilize a 16S approach, which precludes detection of genomics features such as AMPs. Therefore, further research using a shotgun metagenomic approach is needed to understand what role pre-partum teat apex microbiome dynamics play in IMI risk.

## INTRODUCTION

Bovine mastitis is an inflammatory disease of the udder caused by microorganisms that overcome the physical and immunological barriers of the mammary gland (Sordillo et al., 1997). Despite more than a century’s worth of research into the etiology of this disease, mastitis remains an issue for dairy cows and producers (Ruegg, 2017). One major pathogen of concern is *Staphylococcus aureus*, a contagious gram-positive bacteria that results in persistent infections of the mammary gland, and that is notoriously difficult to treat – even with the use of antibiotics (Rainard et al., 2018). This makes bovine mastitis caused by *S. aureus* an important animal health and welfare issue for dairy cows, and a major financial burden for dairy farmers; therefore, further research is needed to prevent and treat these infections (Ruegg, 2017; Rainard et al., 2018).

Perhaps one of the most effective approaches to preventing *S. aureus* intramammary infections (IMIs) is to maintain a low prevalence of the pathogen within the herd by implementing strict biosecurity protocols (Barkema et al., 2009). To achieve this objective, common practices include culling infected animals, maintaining high udder hygiene, separating infected from non-infected animals, and avoiding the purchase of infected animals (Schreiner and Ruegg, 2003; Barkema et al., 2006, 2009). Preventive and treatment practices such as use of antibiotics and internal teat sealants also provide an effective means for treating existing infections and preventing new ones during the dry period, i.e., when the mammary gland is not producing milk (Kabera et al., 2021; McCubbin et al., 2022). However, both antibiotics and internal teat sealants cannot be used in dairy cows from organic-certified milk production systems (Ruegg, 2009).

Indeed, restrictions on antibiotic use have been proposed as a contributing factor to the higher prevalence of *S. aureus* infections observed in some organic farms compared to conventional farms (Pol and Ruegg, 2007a; Ruegg, 2009). This is especially problematic for nulliparous dairy cows because they often calve with *S. aureus* IMIs (De Vliegher et al., 2012). More broadly, the lack of effective measures to treat mastitis on organic farms has led to the use of alternative therapies such as feeding whey products, multivitamins and probiotic therapies, but their impact on mastitis remains unclear (Pol and Ruegg, 2007b; Ruegg, 2009). Even within conventional dairy farms (i.e., those not organically certified), control of *S. aureus* remains a challenge in some circumstances and is considered a major contributor to antimicrobial use (Rainard et al., 2018). Therefore, for both conventional and organic dairy production, research into new tools and therapies for *S. aureus* IMI is critical.

One possible tool that has recently garnered significant attention is the teat apex microbiome (Rainard, 2017), with several studies reporting a potential association with mastitis risk (De Vliegher, 2003; Braem et al., 2014; Falentin et al., 2016). Using culture-independent methods, one study showed that mammary quarters with a history of mastitis had lower microbial diversity compared to teat canals without a history of mastitis (Falentin et al., 2016). Higher microbial richness has also been observed on the teat apices of cows free of an IMI compared to those with an IMI (Andrews et al., 2019). Culture-dependent methods have been used to identify associations between specific microbes on the teat apex and mastitis risk and somatic cell count numbers (De Vliegher, 2003). The presence of non-*aureus* staphylococci (NAS) such as *S. chromogenes* on the teat apex during prepartum has been associated with lower somatic cell counts shortly after calving, which is used as an indicator for subclinical mastitis (De Vliegher, 2003). *S. chromogenes* isolated from the teat apex has also been shown to partially or fully inhibit *S. aureus in vitro* (Ferronatto et al., 2019). The reason for this apparent inhibition is thought to be the production of antimicrobial peptides (AMPs) capable of targeting and attenuating growth of other microorganisms (Cotter et al., 2005). Indeed, NAS isolated from teat apices (including *S. chromogenes*, *S. equorum* and *S. saprophyticus*) have been shown to produce AMPs, resulting in growth inhibition of multiple *S. aureus* strains *in vitro* (Braem et al., 2014). Due to this growing body of literature, the teat apex microbiome and AMPs have been recognized as important areas of further research with regards to mastitis risk (Braem et al., 2014; Rainard, 2017; Rainard et al., 2018); however, few high-throughput sequencing studies of the teat apex microbiome exist, precluding the identification of other microorganisms or AMPs that may be involved in the complex etiology of mastitis.

The primary objective of this study was to identify associations between microbial biomarkers on the teat apex and risk of *S. aureus* IMI in the first 5 weeks after calving in primiparous cows. Secondary objectives were to explore the functional capacity of the teat apex microbiome to produce AMPs, and to identify AMPs that may be associated with *S. aureus* IMI risk.

## MATERIALS AND METHODS

### Study Design

In this case-control study, we collected skin samples from the distal third of the teat apex and quarter milk samples from 710 organically reared dairy cows from 5 farms in Colorado, Minnesota, New Mexico and Texas. Eligibility for farm and cow enrollment has been described previously (Dean et al., 2022). This study was approved by the University of Minnesota Institutional Animal Care and Use Committee (#1807-36109A); the Colorado State University Animal Care and Use Committee (#1402); and the Texas Tech University Animal Care and Use Committee (#18068-10). Collection of skin samples began eight weeks prior to parturition and ended four to five weeks after parturition. Weekly sample collection of quarter milk samples began immediately after parturition and ended four to five weeks after parturition.

### Sample Collection

Skin samples from the distal third of the teat apex were collected in the milking parlor for both prepartum and postpartum animals. Teat disinfectants were not applied by farm personnel prior to sampling. University personnel collected skin samples from each animal by adhering to the following protocol: when working with a new animal, a new pair of gloves was put on; if manure contaminated any teats, then the teats were cleaned with a paper towel; a pre-moistened gauze square was removed from a whirl-pak bag; the gauze square was scrubbed against each of the cow’s teat ends; placed back inside the original whirl-pak bag, tightly sealed and labelled with the cow identifier, date of sample collection and farm name; and then moved into a Ziplock bag that was stored in a cooler with ice. Samples collected in Minnesota were driven back to the University of Minnesota and stored in a freezer at a temperature of -80 °C within 8 hours of sample collection. Samples collected outside of Minnesota were driven back to the participating university and placed in a freezer at a temperature of -80 °C within 8 hours of sampling. Ziplock bags containing samples from outside of Minnesota were then placed inside cryoboxes, filled with dry ice, and shipped overnight to the University of Minnesota for long-term storage at a temperature of -80 °C.

Quarter milk samples from postpartum cows were collected in the milking parlor. University personnel collected quarter milk samples from each animal by adhering to the following protocol: when working with a new animal, a new pair of gloves was put on; for each quarter, milk was forestripped and the ends of each quarter disinfected with a gauze square soaked in 70% EtOH; a flip top tube was then placed under the quarter and approximately 10 mL of milk dispensed into it; the top of the tube was closed and labelled with the quarter that was sampled (e.g., front-left (FL), front-right (FR), rear-left (RL), or rear-right (RR)); and then placed on a metal rack that was moved into a cooler with ice once sampling was completed. Coolers with milk samples collected from Minnesota farms were driven back to the University of Minnesota and stored in a freezer at a temperature of -20 °C within 8 hours. Coolers with milk samples collected from outside of Minnesota were driven back to the participating university and stored in a freezer at a temperature of -20 °C within 8 hours. Milk samples collected from outside of Minnesota were then placed inside cryoboxes, filled with ice packs, and shipped overnight to the University of Minnesota for long-term storage at a temperature of -20 °C.

### Bacterial Culture

Quarter milk samples from each animal and sample collection timepoint were aseptically pooled into composite samples and submitted to the Laboratory for Udder Health at the University of Minnesota for bacterial culture and taxonomic identification (Dean et al., 2022). Pooled milk samples were considered positive for bacterial growth when the number of colony forming units was greater than or equal to one (10 CFU/mL). Pooled milk samples with growth of more than three unique microorganisms were classified as contaminated (Dean et al., 2022). Taxonomic identification of non-contaminated bacterial cultures was carried out using matrix assisted laser desorption ionization-time of flight mass spectrometry (MALDI-TOF MS) (Jahan et al., 2021).

### Participants

Eligibility criteria for being considered a case was based on the presence of *S. aureus* in the postpartum milk samples based on bacterial culture and MALDI-TOF MS. Colonies from milk samples that could only be taxonomically assigned to the genus level (i.e., *Staphylococcus*) were not eligible to be cases, nor were cows with missing calving dates. Eligibility criteria for being considered a control was based on the absence of *S. aureus* in any of the postpartum milk samples based on bacterial culture and MALDI-TOF MS results. Cases and controls were individually matched by farm and days in milk (± 3 days in milk). If a case animal had multiple postpartum milk samples testing positive for *S. aureus*, only the first positive sample was used for matching. If more than one control could be matched to a case animal, then one was selected at random using a random number generator. Cows with a milk sample classified as “contaminated” were not eligible to be considered a case in subsequent weeks. Similarly, cows were not eligible to be matched to cases if a pooled milk sample had been classified as “contaminated” in any previous postpartum milk sample. These decisions were made because contaminated milk samples were not submitted for taxonomic identification using MALDI-TOF; therefore, the presence or absence of *S. aureus* was unknown. Skin samples from matched pairs collected up to and prior to the time of matching were subjected to DNA extraction, library preparation and sequencing.

### Sample Preparation

Prior to DNA extraction, batches of 12 whirl-pak bags containing gauze squares were removed from -80 °C storage and thawed at room temperature. During thawing, a biosafety cabinet was cleaned with 70% EtOH and a glass-bead sterilizer preheated to 250 °C. Once gauze squares were thawed, whirl-pak bags were moved inside the biosafety cabinet. Each whirl-pak bag was processed in the following way: the bag was opened and the gauze square cut into thirds using metal scissors sterilized with a glass-bead sterilizer; the dirtiest third of the gauze square was placed inside a PowerBead Pro Tube using metal clamps sterilized with a glass-bead sterilizer (Qiagen, Cat No. 9002864, Hilden, Germany); the whirl-pak bag was then closed. Once each PowerBead Pro Tube was filled, 800 uL of CD1 lysis buffer was dispensed into each (Qiagen, Cat No. 9002864, Hilden, Germany). PowerBead Pro Tubes were then placed inside a bead- beater for mechanical cell disruption using the following parameters: 20 seconds shake at 2200 revolutions per minute (rpm) and then 30 seconds idle (BioSpec Products, Cat. No. 1001, Bartlesville, OK). This process was repeated three times. PowerBead Pro tubes were then subjected to centrifugation for two minutes at 16,000 RCF. PowerBead Pro tubes were then moved inside a fume hood that was sterilized with 70% EtOH. For each PowerBead Pro tube, 600 uL of the supernatant was dispensed into separate rotor adapters (Qiagen, Cat No. 9002864, Hilden, Germany). Once complete, rotor adapters were placed inside a QIAcube Connect instrument for DNA extraction (Qiagen, Cat No. 9002864, Hilden, Germany).

### DNA Extraction, Library Preparation and Sequencing

DNA was extracted from gauze samples collected form the teat apex using the PowerSoil Pro Kit (Qiagen, Cat No. 47016, Hilden, Germany) on the QIAcube Connect instrument (Qiagen, Cat No. 9002864, Hilden, Germany). Extracted DNA was enzymatically sheared and dual-indexed using the Illumina Nextera XT DNA Library Preparation Kit (Illumina Inc., San Diego, CA). Libraries were created using a quarter (12.5 ul) of the recommended reaction volume (50 ul) of the transposase enzyme and without a AMPure XP Bead Cleanup step, which was shown to generate data with comparable quality and quantity as compared to a full reaction with or without a bead cleanup (**Supplementary File 1; Fig. S1-5**). DNA concentration was measured using a Nanodrop Spectrophotometer (ThermoFisher Scientific). Prior to full-depth sequencing, three separate library pools (“batches”) underwent MiSeq Nano (Illumina Inc., San Diego, CA) quality control sequencing to ensure an equal balance of barcodes across pools (N = 340, 340 and 169 samples in pools 1, 2 and 3, respectively). Once barcode balance was confirmed, each pool was spread across four lanes of an Illumina NovaSeq 6000 S4 flow cell (Illumina Inc., San Diego, CA). Three positive control samples were included in each pool (ZymoBIOMICS, Cat No. D6310). Batch effects by lane were minimized by spreading libraries across each of the four lanes of the S4 flow cell. Batch effects by sequencer were minimized by randomizing samples onto separate sequencing runs. In other words, samples from cases were not sequenced on one sequencing run and controls sequenced on another. Doing so would have completely confounded our independent variable – case or control group – and precluded us from determining any relationship between the teat apex microbiome and intramammary infections caused by *S. aureus*.

### Bioinformatics

The Nextflow pipeline framework was used for all data processing of sequence data (Di Tommaso et al., 2017).

### Quality Control

Paired-end sequence reads were inspected for data quality and adapter contamination using FastQC (https://github.com/s-andrews/FastQC) and MultiQC (Ewels et al., 2016). Sequence reads were subjected to quality trimming using TrimGalore (Krueger et al., 2021): one-base pair was trimmed from the 3’ ends of each sequence read; and then subsequences with a quality score less than 25 or at least a four base-pair overlap with the Nextera adapter sequence were trimmed (‘CTGTCTCTTATA’). After trimming, sequence reads were removed if they contained any ambiguous base pairs; had a sequence length less than 50 base pairs; or were missing their mate pair. Only sequence reads that survived this QC process were used in downstream analyses.

### Background DNA Removal

Sequence reads were aligned to a custom reference library to identify bovine sequence reads. The custom library was built from the *Bos taurus* (RefSeq assembly accession: GCF_002263795.2) reference genome using Kraken2’s ‘download-taxonomy’, ‘add-to-library’ and ‘build’ commands (Wood et al., 2019). Sequence reads that were not classified as bovine were used as input for taxonomic assignment.

### Taxonomic Assignment

Sequence reads were aligned to the CHOCOPhlan marker-gene database using Metaphlan with the ‘add_viruses’ parameter (Beghini et al., 2021). The queries included alignments to archaea, bacteria, eukaryotes, and viruses. We noted that some of the taxonomic assignments inferred by Metaphlan were likely incorrect for some virus classifications; therefore, we chose to describe these viruses at a broader taxonomic rank in the text and within graphs when appropriate (e.g., at the order level instead of the species level).

### Identification of Antibacterial Peptides

The Data Repository of Antimicrobial Peptides (DRAMP 3.0) was selected as the reference database for AMP characterization (Shi et al., 2022). Non-host, paired-end sequence reads were merged using the ‘read_merger’ program implemented in Kraken (Wood and Salzberg, 2014).

Merged sequence reads were aligned to the DRAMP protein database using PALADIN (Westbrook et al., 2017). The ‘index’ and ‘align’ commands in PALADIN were used in series to index the DRAMP database and then align genomic sequences to the DRAMP database.

Sequence alignment map (SAM) files output from PALADIN were converted to sorted binary alignment map (BAM) files using SAMtools (Li et al., 2009). Using the BAM file, a count matrix was generated by counting the number of sequence reads in each sample that aligned to each reference sequence in the DRAMP database using seqkit (Shen et al., 2016) (bit 4 unset). Only primary alignments were considered for downstream analysis. Reads with multiple hits to the reference database (indicated by the XA tag in the SAM file) were retained. Alignments to peptides labelled as “synthetic” in the “protein existence” column of the DRAMP reference database were discarded, as were those with a eukaryotic origin.

### Analysis of Positive Controls

Reference genomes for the positive mock community were downloaded from https://s3.amazonaws.com/zymo-files/BioPool/ZymoBIOMICS.STD.refseq.v2.zip

(ZymoBIOMICS, Cat No. D6310). Sequence reads from 10 positive control samples were aligned to the Zymo reference database using BWA-MEM (Li, 2013). The resulting SAM file was converted to a sorted BAM file using SAMtools (Li et al., 2009). Counts of reads aligning to each reference sequence – and associated plasmids – were tallied from each BAM file using the seqkit program with the ‘bam’ option and ‘count’ flag (Shen et al., 2016). Only counts of primary alignments were considered. Counts from each sample were converted to relative abundances and visualized using stacked bar charts (Wickham, 2009).

### Sequencing Statistics

Linear mixed effect models were used to determine if the raw sequencing depth (primary outcome) of teat apex samples differed between cases and controls after adjusting for potential confounding variables, such as sequencing batch, weeks to infection, days to infection and farm (Bates et al., 2015). Confounding factors remained in the model if their effect was significantly associated with the primary outcome. This was determined by comparing a full model -- with each of the above variables -- to a null model with one of the variables removed using a likelihood ratio test. The same procedure was used to determine if the host sequencing depth (secondary outcome) differed between cases and controls. The ‘emmeans’ package was used to calculate adjusted raw or host sequencing depths between cases and controls when significantly associated with the outcome (Lenth et al., 2022). Univariate linear regression and ordinal logistic regression were used to determine if days in milk, days to infection or weeks to infection differed between cases and controls (Ripley et al., 2023). A permutational multivariate analysis of variance was used to determine if sequencing batch was significantly associated with teat apex microbiome beta-diversity (Anderson, 2001; Oksanen et al., 2022).

### Microbiome Statistics

Logistic regression was used to determine the association between the presence of a *S. aureus* IMI (as determined from the milk sample cultures) and the odds of observing each microorganism in the metagenomic data generated from the teat apex samples. The term operational taxonomic unit (OTU) will be used throughout the text to refer to the microbial features (i.e., microorganisms) inferred by Metaphlan. Logistic models were fit for each OTU using the *glmer* function in the ‘lme4’ package (Bates et al., 2015). For each OTU, a logistic model with a ‘logit’ link function and ‘binomial’ family was fit using notation from (Dohoo et al., 2009):

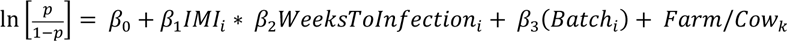

where *p* represents the probability of observing the OTU; ln(p/(1-p)) represents the logit transformation; *IMI* is a two-level factor representing the presence or absence of a *S. aureus* IMI; *WeeksToInfection* is a seven-level factor representing the number of weeks prior to IMI at which the sample was taken; *Batch* is a three-level factor representing sequencing batch; *Farm* is a five-level factor representing the farm the sample was collected from; and *Cow* is a multi-level factor representing the cow that was sampled. A nested random effect was input into the model to account for cows nested within each farm. The ‘emmeans’ package was used to calculate adjusted risks and odds ratios for each OTU on the response scale (Lenth et al., 2022). Adjusted risks were defined as the proportion of samples with the presence of the OTU after accounting for other covariates in the model. Confidence intervals were used to determine if the odds of harboring a given OTU were significantly higher in cases or controls; therefore, *P*-values were not subjected to adjustment for multiple comparisons.

Negative binomial regression models (NB) were used to investigate the association between the presence of a *S. aureus* IMI and the abundance of each OTU in cases and controls. OTUs present in fewer than 5% of samples were discarded. The negative binomial framework was chosen due to the overdispersion observed in our dataset (**Supplementary File 2, Fig. S1**).

Indeed, these are common characteristics of high-throughput sequencing experiments (Fang et al., 2014; Love et al., 2014; Zhang et al., 2018; Zhang and Yi, 2020) and numerous software packages exist to account for these very features (Mallick et al., 2021; Fernandes et al., 2014; Love et al., 2014; Paulson et al., 2013). However, few provide convenient and robust methods for handling datasets produced from observational studies, e.g., testing for interactions, specifying multiple (or nested) random effects; and dealing with repeated measurements; therefore, we chose to use the models implemented in the glmmTMB package (Brooks et al., 2017) because they offer these features and integrate well with downstream statistical packages such as emmeans (Lenth et al., 2022). Negative binomial regression models were fit for each OTU in the following way:

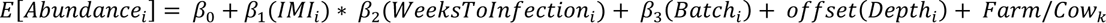

An offset term for sequencing depth was included in the model to account for differences in sequencing depth per sample, as in (Fang et al., 2014). Our response variable for each statistical model (i.e., abundance) was not normalized a priori simply because the negative binomial model expects counts and not fractional or negative integers, which are commonly produced by many normalization methods and transformations (cumulative sum scaling (Paulson et al., 2013), base 10 logarithm or centered-log ratio (Aitchison, 1982)). The significance of the interaction term was tested using a likelihood ratio test. If the term did not significantly improve model fit, then the term was removed, and the additive model ran instead. Significance values were corrected for multiple comparisons using Benjamin Hochberg correction (Benjamini and Hochberg, 1995). Fits from each model were used as input to the emmeans function in the ‘emmeans’ package to calculate adjusted OTU abundances. The adjusted abundances from cases and controls were used to calculate a log fold change for each OTU. Intraclass correlation coefficients were computed for variance components – when possible – using the *icc* function in the ‘performance’ package (Lüdecke et al., 2021).

Linear mixed effect models were used to investigate the association between the presence of an*S. aureus* IMI and antimicrobial peptide (AMP) richness and diversity. AMP richness and diversity were measured using the *estimate_richness* function in ‘phyloseq’ (McMurdie and Holmes, 2013). For each alpha diversity outcome (richness or diversity), the following linear model was fit:

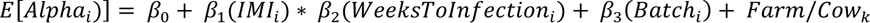

The ‘emmeans’ package was used to estimate adjusted richness and diversity values between cases and controls for each alpha diversity measure on the response scale (Lenth et al., 2022). AMP abundances were compared between cases and controls using the same method and formula described for the comparison of OTU abundances.

## RESULTS

### Animal and Herd Characteristics

Animal and herd characteristics for each of the five enrolled farms in this study are described in **Table 1**. The predominant animal breed for four of the five farms was Holstein, and a mix of Holstein crosses for one of the five farms. The number of milking cows for enrolled farms ranged from 100 to 3,000. For four of the five farms, DHI testing was confirmed by farm personnel; quarter milk culture was performed on quarters with clinical mastitis; and use of fly control strategies was practiced. Pre- and post-milking teat disinfectants were used on all farms.

**Table 1.**
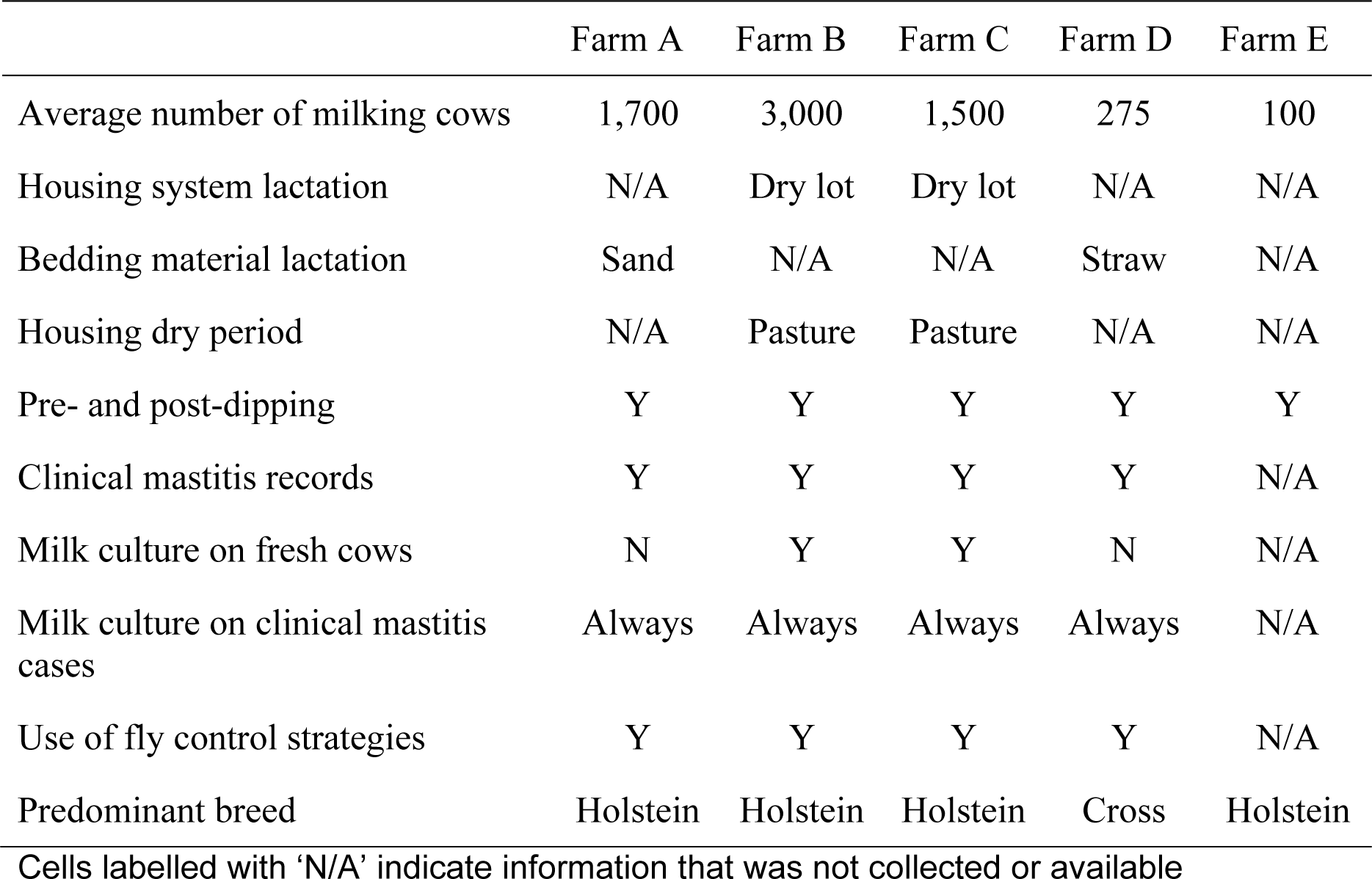
Herd and management information of enrolled farms.

### Enrollment and Sample Collection Information

The number of animals enrolled in this study and the number of samples collected from participating farms is described in **Table 2**. A total of 710 animals were enrolled from five farms, with a range of between 28 and 288 animals enrolled on each farm. The total number of teat apex samples collected from the 710 animals was 4,827. The median number of teat apex samples collected from each animal was 8. Samples were collected in 2019 and parts of 2020.

**Table 2.**
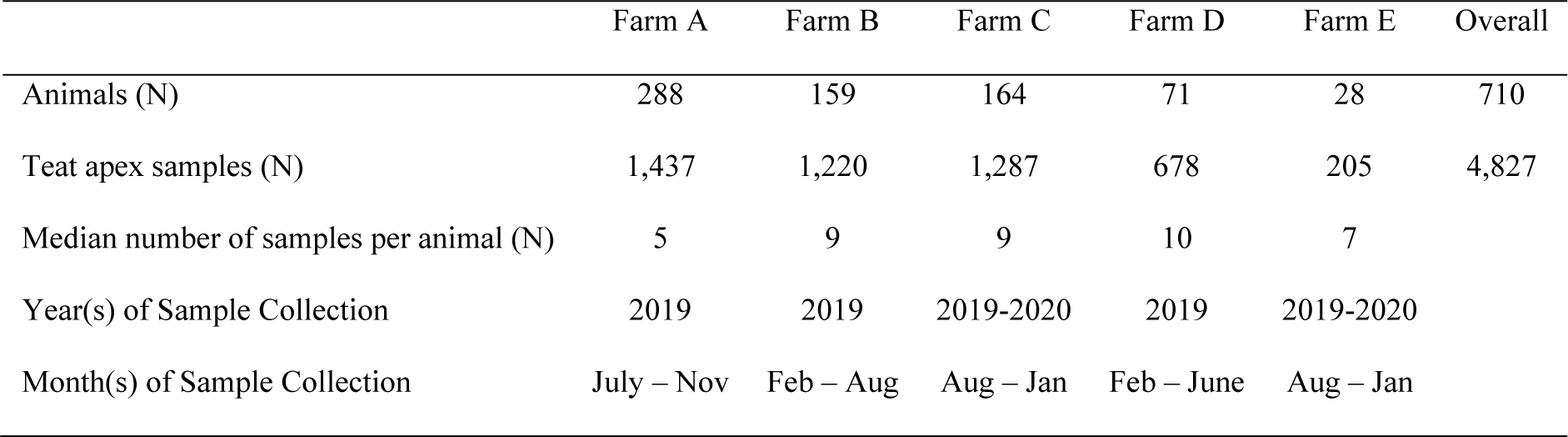
Animal enrollment and sample collection information.

|  | Farm A | Farm B | Farm C | Farm D | Farm E | Overall |
| --- | --- | --- | --- | --- | --- | --- |
| Animals (N) | 288 | 159 | 164 | 71 | 28 | 710 |
| Teat apex samples (N) | 1,437 | 1,220 | 1,287 | 678 | 205 | 4,827 |
| Median number of samples per animal (N) | 5 | 9 | 9 | 10 | 7 |  |
| Year(s) of Sample Collection | 2019 | 2019 | 2019-2020 | 2019 | 2019-2020 |  |
| Month(s) of Sample Collection | July – Nov | Feb – Aug | Aug – Jan | Feb – June | Aug – Jan |  |

### Case-Control Matching Statistics

The number of animals selected for this case-control study and their sample characteristics are described in **Table 3**. One of the five farms - Farm E - did not have any cases of *S. aureus* IMIs; therefore, no samples from this farm were selected for this case-control study. Based on our eligibility criteria, the total number of cows included in this case-control study was 162 (cases = 81, controls = 81). A total number of 839 teat apex samples from these 162 cows were available for metagenomic sequencing (418 from cases and 421 from controls). The median number of samples sequenced from cases was 5 (min = 3; max = 9) and controls was 5 (min = 3; max = 8). The median number of days after calving for when a case became a case was 6 (min = 0, max = 32). The median number of days after calving for when a control was matched to a case was 6 (min = 0, max = 29).

**Table 3.** Case-control matching characteristics.

|  | Case | Control | Total |
| --- | --- | --- | --- |
| No. of animals | 81 | 81 | 162 |
| No. of samples | 418 | 421 | 839 |
| No. of animals by farm |  |  |  |
| Farm A | 20 | 20 |  |
| Farm B | 31 | 31 |  |
| Farm C | 9 | 9 |  |
| Farm D | 21 | 21 |  |
| Farm E | 0 | 0 |  |
| No. of Samples Per Animal (Min, Max, Median) | 3,9,5 | 3,8,5 |  |
| Median DIM at time of matching | 6 | 6 |  |

### Sequencing Statistics

A total number of 26.5B (billion) sequence reads was generated from 839 teat apex samples sequenced on three separate runs of an Illumina NovaSeq using an S4 flow cell (**Supplementary Table 1**) (Mean = 31.6M reads per sample; SD = 6.0M reads). After QC, 25.7B sequence reads remained (Mean = 30.6M reads per sample; SD = 5.8M reads); and after removal of *Bos taurus* reads, 18.0B sequence reads remained (Mean = 21.5M reads per sample; SD = 9.9M reads).

Sequencing batch, case or control group, farm, days to infection and weeks to infection were not significantly associated with raw sequencing depth (**Fig. 1 A-E**, ANOVA *P* > 0.05). Sequencing batch was not significantly associated with host sequencing depth (**Fig. 1F**; i.e., *Bos taurus* DNA), but case or control group, farm and weeks to infection were (ANOVA *P* < 0.05). On average, cases had 4.25 million more *Bos taurus* DNA reads compared to controls after adjusting for weeks to infection and farm (95% CI: 2.62M – 5.87M; *P* < 0.0001; **Fig. 1 G-J**).

**Fig. 1.**
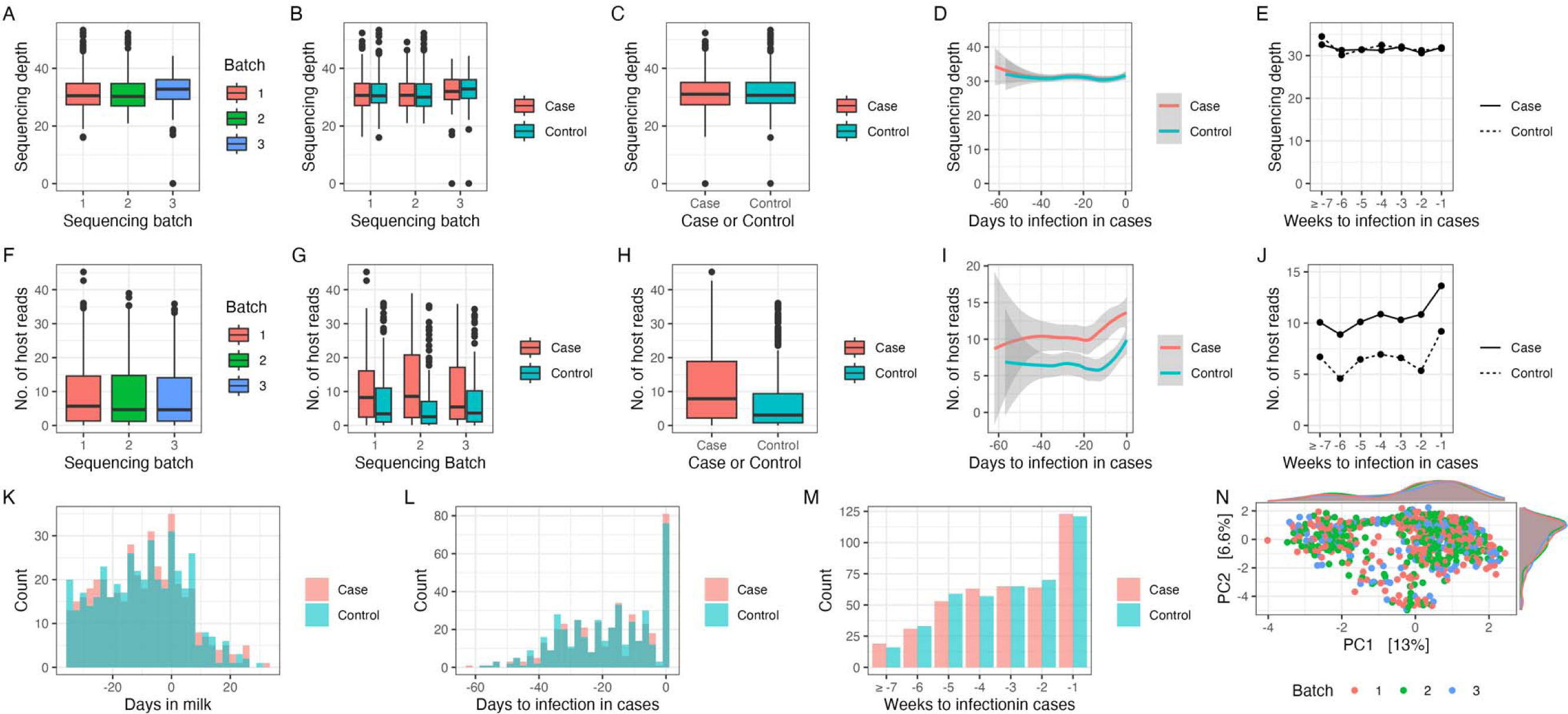
Sequencing and quality control plots of teat apex samples. (A) Raw sequencing depth as a function of sequencing batch; (B) Raw sequencing depth as a function of case or control group and sequencing batch; (C) Raw sequencing depth as a function of case or control group; (D) Raw sequencing depth as a function of case or control group and days to infection; (E) Raw sequencing depth as a function of case or control group and weeks to infection; (F) Host sequencing depth as a function of sequencing batch; (G) Host sequencing depth as a function of case or control group and sequencing batch; (H) Host sequencing depth as a function of case or control group; (I) Host sequencing depth as a function of case or control group and days to infection; (J) Host sequencing depth as a function of case or control group and weeks to infection; (K) Count of the number of teat apex samples by case or control group and days in milk; (L) Count of the number of teat apex samples by case or control group and days to infection; (M) Count of the number of teat apex samples by case or control group and weeks to infection; and (N) PCA plot of teat apex microbiome colored by sequencing batch.

Days in milk, days to infection and weeks to infection in cases did not differ between cases and controls (**Fig. 1 K-M**). Clustering of the teat apex microbiome by sequencing batch was not visually apparent using PCA (**Fig. 1N**); indeed, the amount of variation in the teat apex microbiome that could be explained by sequencing batch was < 1% (*R*^2^ = 0.0036, F = 1.53; *P* = 0.0009).

### Positive Control Statistics

Ten positive control samples (i.e., mock communities) were sequenced on three separate runs of the Illumina NovaSeq instrument (Illumina Inc., San Diego, CA). The composition of the mock community included 8 bacteria: *Listeria monocytogenes*, *Pseudomonas aeruginosa*, *Bacillus subtilis*, *Escherichia coli*, *Salmonella enterica*, *Lactobacillus fermentum*, *Enterococcus faecalis* and *Staphylococcus aureus*; and 2 eukaryotes: *Cryptococcus neoformans* and *Saccharmomyces cerevisiae* (Zymo Research Cat. No. D6310). All 10 members of the mock community were identified from 100% of positive control samples (**Supplementary File 2 Fig. S2**); however, the relative abundance of these mock community members in our data differed from the theoretical composition. The most abundant member of the mock community (*L. monocytogenes*) was under-represented with a mean relative abundance of 60.9% (theoretical: 89.1%). All other members of the mock community tended to be over-represented compared to their theoretical compositions (**Supplementary File 2 Table S1**).

### Taxonomic Classification Statistics

A list of all microorganisms identified from shotgun metagenomic sequencing of the teat apex microbiome are listed in **Supplementary Table 2**. Non-host sequencing reads were taxonomically assigned to 1,395 archaeal, bacterial, eukaryotic, and viral species. The number of distinct species classified to each kingdom was 18 for archaeal species (1.3% of all 1,395 identified OTUs), 987 for bacterial species (70.8%), 10 for eukaryotic species (0.7%), and 380 for viral species (27.2%).

The top 60 most prevalent microorganisms and their abundances from the metagenomic data (crude prevalence from 839 samples) are shown in **Fig. 2**. Prevalent bacteria included NAS: *S. chromogenes* (n = 472 samples, 56.2%), *S. haemolyticus* (n = 470, 56.0%), *S. auricularis* (n = 412, 49.1%) and *S. devriesei* (n = 388, 46.2%); *Corynebacterium* species: *C. efficiens* (n = 732, 87.2%), *C. xerosis* (n = 733 87.3%), *C. marinum* (n = 732, 87.2%); *Bifidobacterium* species: *B. pseudolongum* (n = 827, 98.5%) and *B. merycicum* (n = 548, 65.3%); *Jeotgalicoccus sp* (n = 679, 80.9%); *Dietzia alimentaria* (n = 660, 78.6%); *Kocuria polaris* (n = 610, 72.7%). The most prevalent archaeal species were methane producing bacteria: *Methanosarcina mazei* (n = 276, 32.8%), *Methanobrevibacter thaueri* (n = 261, 31.1%) and *Methanobrevibacter millerae* (n = 125, 14.8%). The most prevalent viral species were two herpes-associated viruses, which we describe at the order-level taxonomy: *Herpesvirales I* (n = 836, 99.6%) and *Herpesvirales II* (n = 346, 41.2%); bacteriophages: *Microbacterium phage Min 1* (n = 579, 69.0%), *Staphylococcus virus EW* (n = 268, 31.9%) and *Enterobacteria phage P4* (n = 267, 31.8%); and a bovine-associated polyomavirus (n = 518, 61.7%). The most prevalent eukaryotic species was the fungus *Aspergillus fumigatus* (n = 13, 1.5%).

**Fig. 2.**
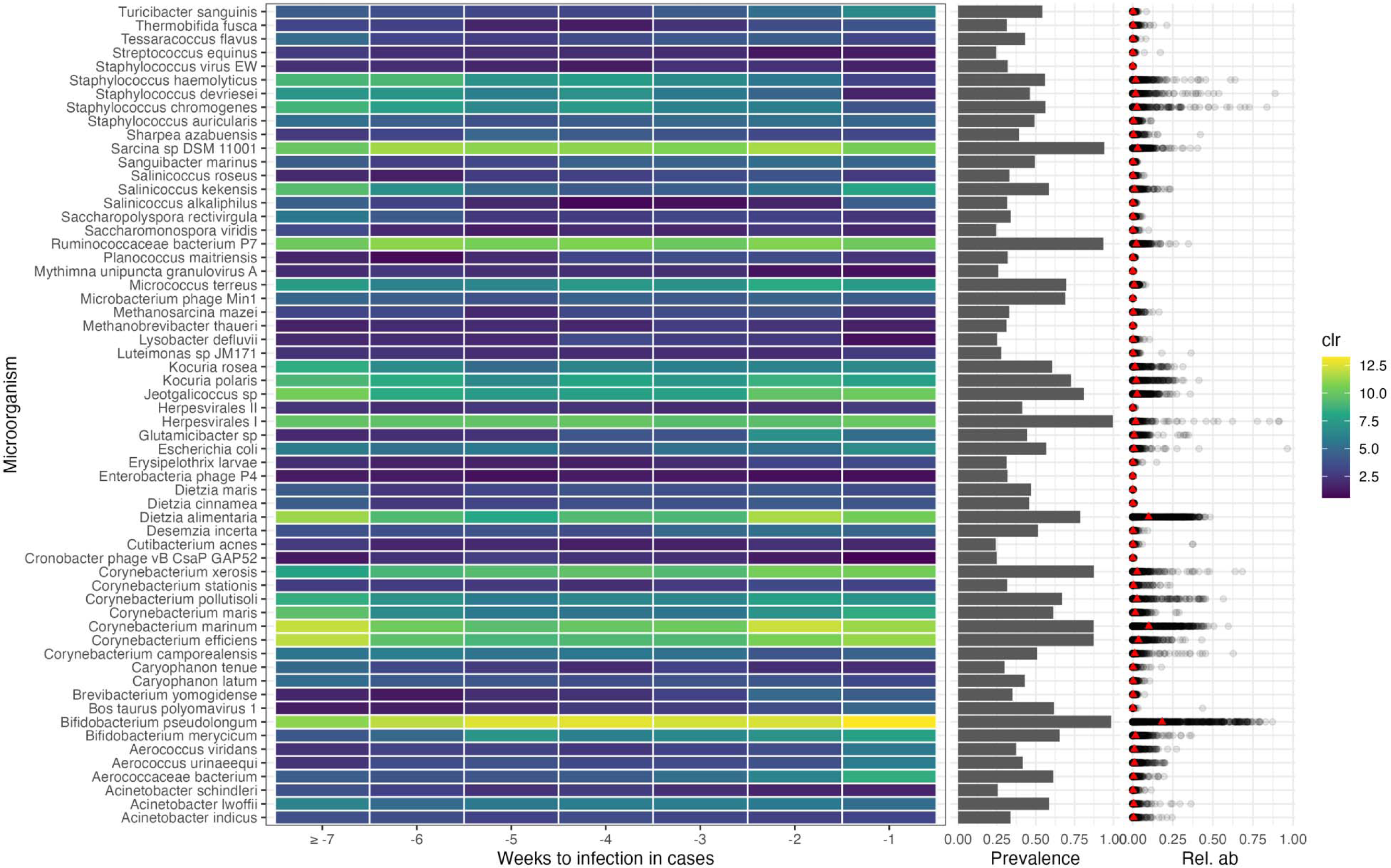
Taxonomic composition, prevalence and abundance of the most frequently isolated microorganisms from the teat apex microbiome. The signature heatmap displays the median normalized abundance (clr: center-log ratio) of each microorganism for each week relative to infection in cases. The bar graph shows the prevalence of microorganisms in teat apex samples (denominator 839 samples). The dot plot shows the relative abundance of each microorganism within each sample. The red triangles represent the mean relative abundance of each microorganism across all samples.

The most abundant microorganisms were bacteria from the NAS, *Corynebacterium*, *Bifidobacterium*, *Dietzia* and *Jeotgalicoccus* genera, accounting for much of the metagenomic DNA from the teat apex (**Fig. 2**). The three most abundant species from the metagenomic data were *B. pseudolongum* (18.3%), *C. marinum* (10.0%) and *D. alimentaria* (9.8%). Among the NAS group, *S. chromogenes (2.5%)*, *S. haemolyticus (1.8%)*, *S. devreisei (1.8%)* were the most abundant. Other abundant *Corynebacterium* species included *C. marinum* (10.0%), *C. efficiens (3.5%)*, *C. xerosis (2.7%)*, and *C. pollutisoli (2.6%)*. The most abundant species within the *Jeotgalicoccus* genera was an un-speciated *Jeotgalicoccus* sp. (2.6%). *Sarcina sp.* (3.0%); *Kocuria polaris* (2.2%) and *K. rosea* (1.3%); and *Ruminococcacaea P7* species (2.2%) were abundant as well.

Presence of *Staphylococcus aureus* and skin-associated viruses on the teat apex prior to parturition were the strongest risk factors for *Staphylococcus aureus* IMIs after parturition The presence of *S. aureus* and skin-associated viruses in the metagenomic teat apex data prepartum were significant risk factors for postpartum *S. aureus* IMI (**Fig. 3A**). Samples from cases had significantly higher odds of harboring *S. aureus* DNA in the teat apex metagenome compared to controls (OR = 38.95, 95% CI: 14.8-102.2). The adjusted risk of *S. aureus* DNA in samples from cases was 33.6% (95% CI: 22.5-46.8%) and controls was 1.2% (95% CI: 0.4- 3.3%). The odds of harboring *S. aureus* DNA varied more between cows (intraclass correlation coefficient (ICC): 0.39) than it did between farms (ICC: 0.01). In some cows, the presence of *S. aureus* DNA on the teat apex could be observed ≥ 7 weeks prior to the detection of the postpartum *S. aureus* IMI (**Fig. 3B**), almost 62 days away from detection of the infection (**Fig. 3C**). A majority of *S. aureus* DNA found on the teat apex was observed prior to parturition, with few samples containing any detected *S. aureus* DNA after parturition (**Fig. 3D**). Cases also had higher odds of containing DNA from a herpes-associated virus -- *Herpesvirales* II virus (OTU1326) -- (OR = 2.09, 95% CI: 1.46-2.98) compared to controls. The adjusted risk of *Herpesvirales* virus II DNA in cases was 43.57% (95% CI: 28.74-59.65%) and controls was 26.97% (95% CI: 16.07-41.59%). The variation in the odds of harboring *Herpesvirales* virus II DNA was similar between cows (ICC: 0.08) and farms (ICC: 0.08). Samples from cases also had a higher odds of containing DNA from an unspeciated *Psychrobacter* bacterium (OR = 1.83, 95% CI: 1.82–1.83), but the prevalence of this bacterium was low (n = 82 total samples) and only found on a single farm.

**Fig. 3.**
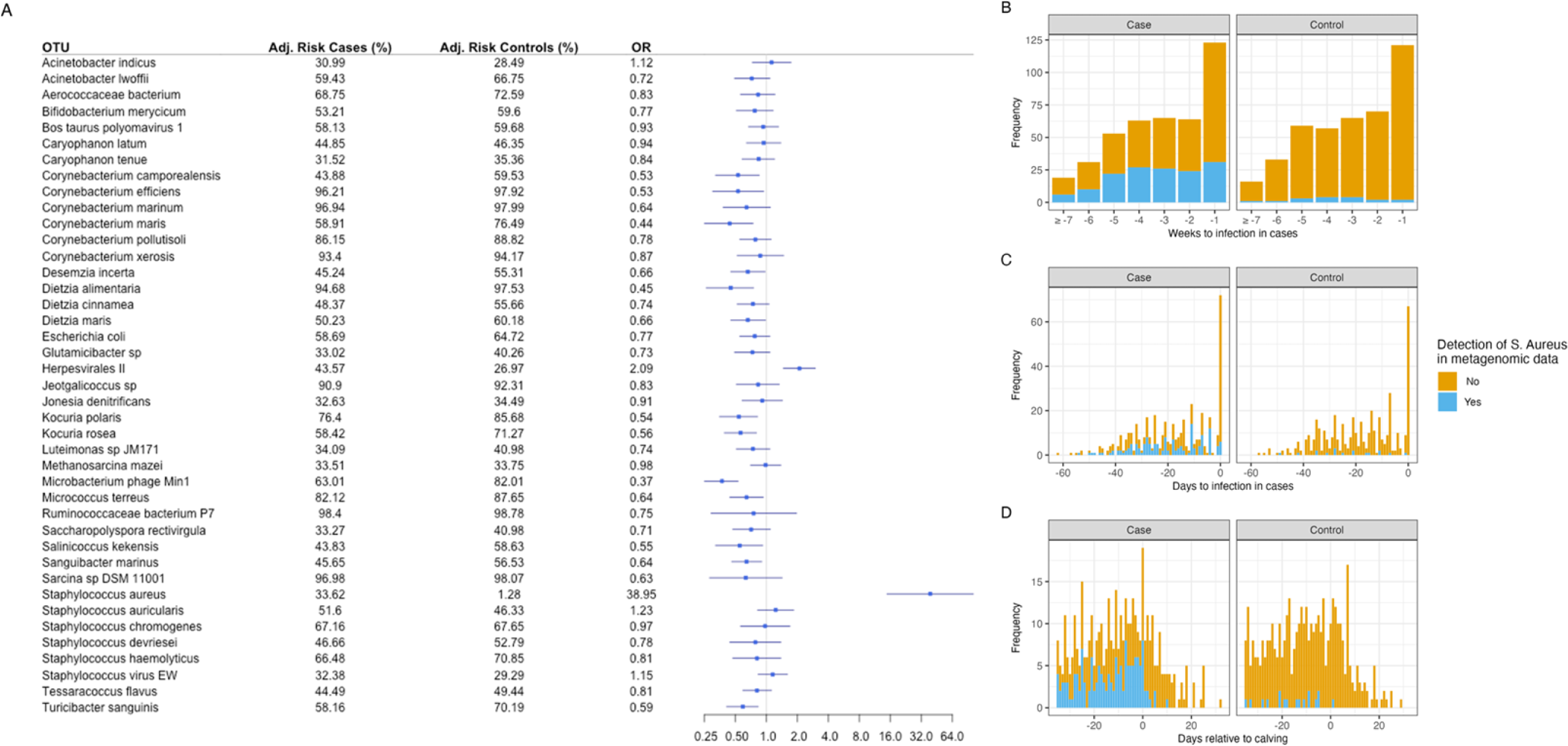
**A.** Forest plot describing the relationship between having an *S. aureus* intramammary infection and the odds of each OTU being present on the teat apex. Adjusted risks (Adj. Risk) represent the probability of the OTU being present on the teat apex, stratified by cases and controls. Odds ratios (OR) represent the odds of the OTU being present in cases relative to controls. Dashed vertical line in the forest plot indicates the null association. Points represent ORs and horizontal bars represent 95% confidence intervals. The number of samples (“Frequency”, y-axes) with *S. aureus* in the teat apex metagenomic data (No = orange, Yes = blue) as a function of Weeks to Infection (**B**); Days to Infection (**C**); and Days Relative to Calving (**D**).

A total number of 31 bacteria and 2 bacteriophages were identified as potential microbial protectives (i.e., OTUs associated with lower odds of *S. aureus* IMI, **Fig. 3A**). The most prevalent microorganisms in controls associated with protection were *Microbacterium* phage Min 1 (OR = 0.37, 95% CI: 0.25-0.53), *Corynebacterium efficiens* (OR = 0.53, 95% CI: 0.30-0.94), *Kocuria polaris* (OR = 0.54, 95% CI: 0.35-0.82), *Micrococcus terreus* (OR = 0.64, 95% CI: 0.44-0.93), *Dietzia alimentaria* (OR = 0.45, 95% CI: 0.26-0.75), *Corynebacterium maris* (OR = 0.44, 95% CI: 0.25-0.74), *Salinicoccus kekensis* (OR = 0.55, 95% CI: 0.32-0.92) *Turicibacter sanguinis* (OR = 0.59, 95% CI: 0.41-0.83) and *Corynebacterium camporealensis* (OR = 0.53, 95% CI: 0.32-0.85).

The odds of NAS such as *Staphylococcus chromogenes* (OR = 0.97, 95% CI: 0.56-1.69), *Staphylococcus devriesei* (OR = 0.78, 95% CI: 0.44-1.38) or *Staphylococcus haemolyticus* (OR = 0.81, 95% CI: 0.47-1.38) being present on the teat apex was not different between cases and controls. The adjusted risk of *Staphylococcus chromogenes* DNA in samples from cases was 67.1% (95% CI: 46.2-82.9) and controls was 67.6% (95% CI: 46.8-83.2). The adjusted risk of *Staphylococcus devriesei* DNA in cases was 46.6% (95% CI: 22.0-73.0) and controls was 52.7% (95% CI: 26.5-77.6). The adjusted risk of *Staphylococcus haemolyticus* DNA in cases was 66.4% (95% CI: 54.2-76.8%) and controls was 70.8% (95% CI: 59.0-80.3).

### Abundance of *Staphylococcus aureus* on the teat apex was higher in cases compared to controls

The abundance of sequence reads originating from *S. aureus* was significantly higher in cases compared to controls (LogFC = 23.8, BH-adjusted *p* < 0.0001, **Fig. 4A**). Across all farms, the abundance of *S. aureus* was consistently higher in cases compared to controls (**Fig. 4B**). There was more variation in *S. aureus* abundance between cows (ICC: 0.66) than farms (ICC: 0.006). The abundance of sequence reads originating from *Psychrobacter immobilis* (LogFC = 10.23, BH-adjusted *p* = 0.002) and *Bacteroides pyogenes* (LogFC 7.34, BH-adjusted *p* = 0.04) was significantly higher in samples from controls compared to cases (**Fig. 4A**). Unlike *S. aureus*, these two bacteria were primarily found on a single farm (**Fig. 4B**), and this was reflected in their variance components for the farm specific random effects for *P. immobilis* (ICC: 0.96) and *B. pyogenes* (ICC: 0.56). The abundance of sequence reads originating from NAS -- *S. chromogenes*, *S. devriesei*, *S. haemolyticus* and *S. auricularis* -- were not significantly different between cases and controls (**Fig. 4A and Fig. 4B**).

**Fig. 4.**
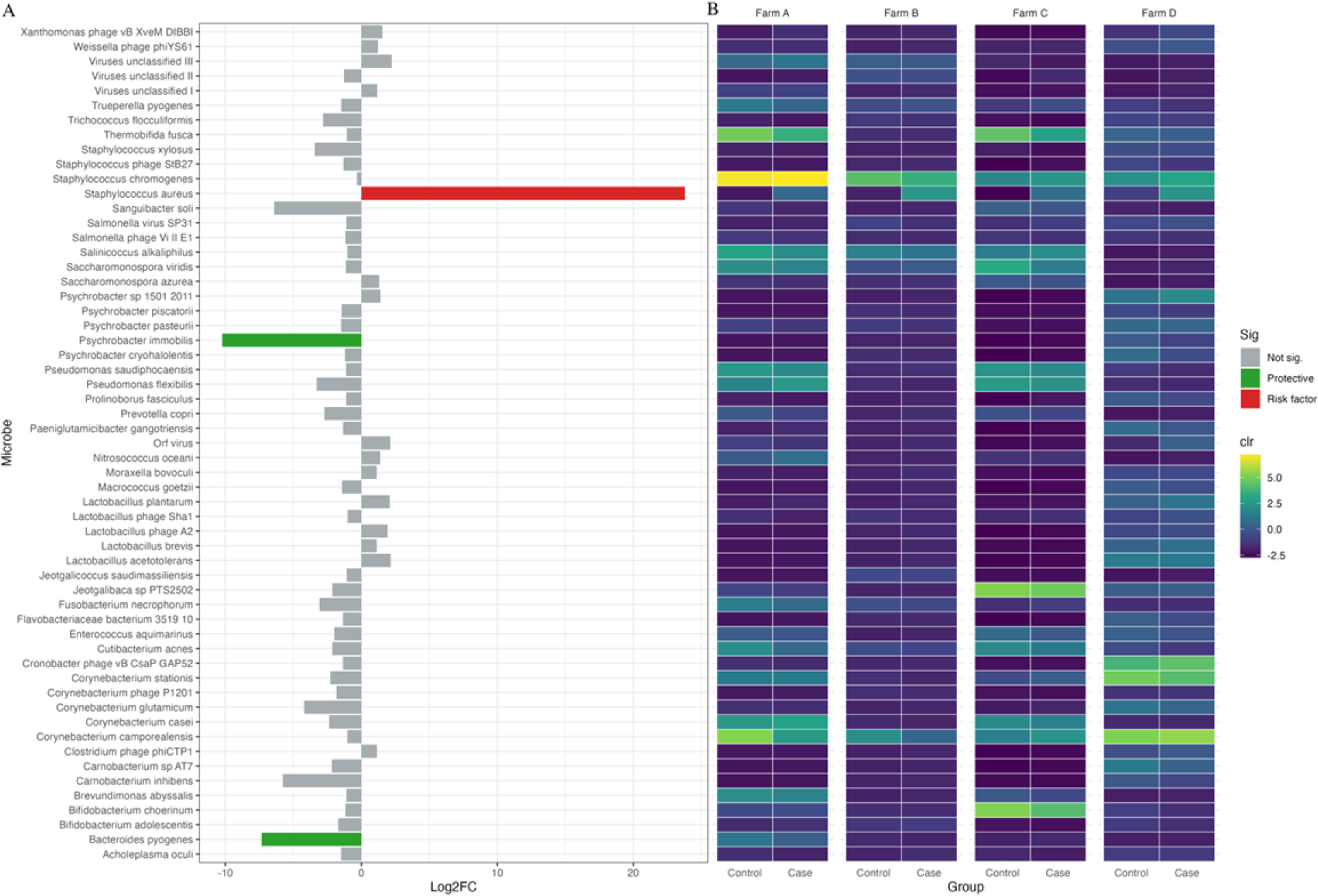
**A.** Bar plot displaying log fold change in abundance (x-axis) of each microorganism (y-axis) between cases and controls. Bars colored red indicate microorganisms with a significantly higher abundance in samples from cases compared to controls (i.e., risk factor); bars colored green indicates microorganisms with a significantly higher abundance in samples from controls compared to cases (i.e., protective); bars colored grey indicate microorganisms without a significant effect size (i.e., not sig). **B.** Heatmap displaying normalized microbial abundances between cases and controls, stratified by farm. Abundances represent centered log ratios (i.e., clr).

### Antimicrobial Peptides

We identified 171 unique putative AMPs across all 839 metagenomic datasets generated from teat apex samples. After removing AMPs with a synthetic (N = 16) and eukaryotic (N=81) origin, 74 AMPs remained for analysis (**Supplementary Table 3**). Two samples – one in the case and one in the control group - did not have any bacterial AMPs. Many of the putative bacterial AMPs that we identified from the teat apex were rare, with 64 of 74 (86.4%) being found in fewer than 10 samples (**Fig. 5**). The mean number of bacterial AMPs found in cases was 1.45 (95% CI: 1.33-1.75) and controls was 1.63 (95% CI: 1.42-1.84). The difference in the mean number of AMPs was not statistically different between cases and controls (contrast: case - control; estimate = -0.09, *p* = 0.13). The mean AMP diversity in cases was 1.09 (95% CI: 1.05- 1.14) and controls was 1.08 (95% CI: 1.03-1.13), with a non-statistically significant mean difference of 0.13 (contrast: cases - control; *p* = 0.36). The most prevalent and abundant predicted bacterial AMPs were Class I (Microcin B17) and IIa microcins (Colicin-V). Microcin B17 was found in 837 of 839 teat apex samples (99.7%) and Microcin-V was found in 43 of 839 teat apex samples (5.1%). The adjusted mean abundance of Microcin B17 in cases was 27.9 (95% CI: 23.5-33.2) and controls was 33.6 (95% CI: 28.3-39.9), with a significant mean difference (*p* = 0.0014). As the remaining putative AMPs identified from the teat apex were rare, we chose not to perform a statistical analysis on them; therefore, crude statistics are reported. Two homology-inferred (DRAMP03172 and DRAMP03170) and three protein-level (DRAMP02795, DRAMP02794, DRAMP02793) AMPs belonging to the staphylococcal hemolytic protein family were found in 108 of 839 samples (12.8%): antibacterial protein 1 homolog in 42 samples (5.0%); antibacterial protein 3 homolog in 62 samples (7.3%); antibacterial protein 1 in one sample (0.1%); antibacterial protein 2 in 28 samples (3.3%) and antibacterial 3 protein in 29 samples (3.4%). The number of samples with any of these AMPs was 42 in cases (5.0%) and 66 in controls (7.8%). A class I bacteriocin named Sonorensin, typically found in *Bacillus sonorensis* MT93 strains, was found in 31 samples (3.6%). Class II bacteriocins typically found in *S. aureus* were found in 22 samples (2.6%): Aureocin A70 (AurA, 9 samples), Aureocin A70 (AurB, 5 samples), Aureocin A70 (AurC, 3 samples), Aureocin A70 (AurD, 5 samples).

**Fig. 5.**
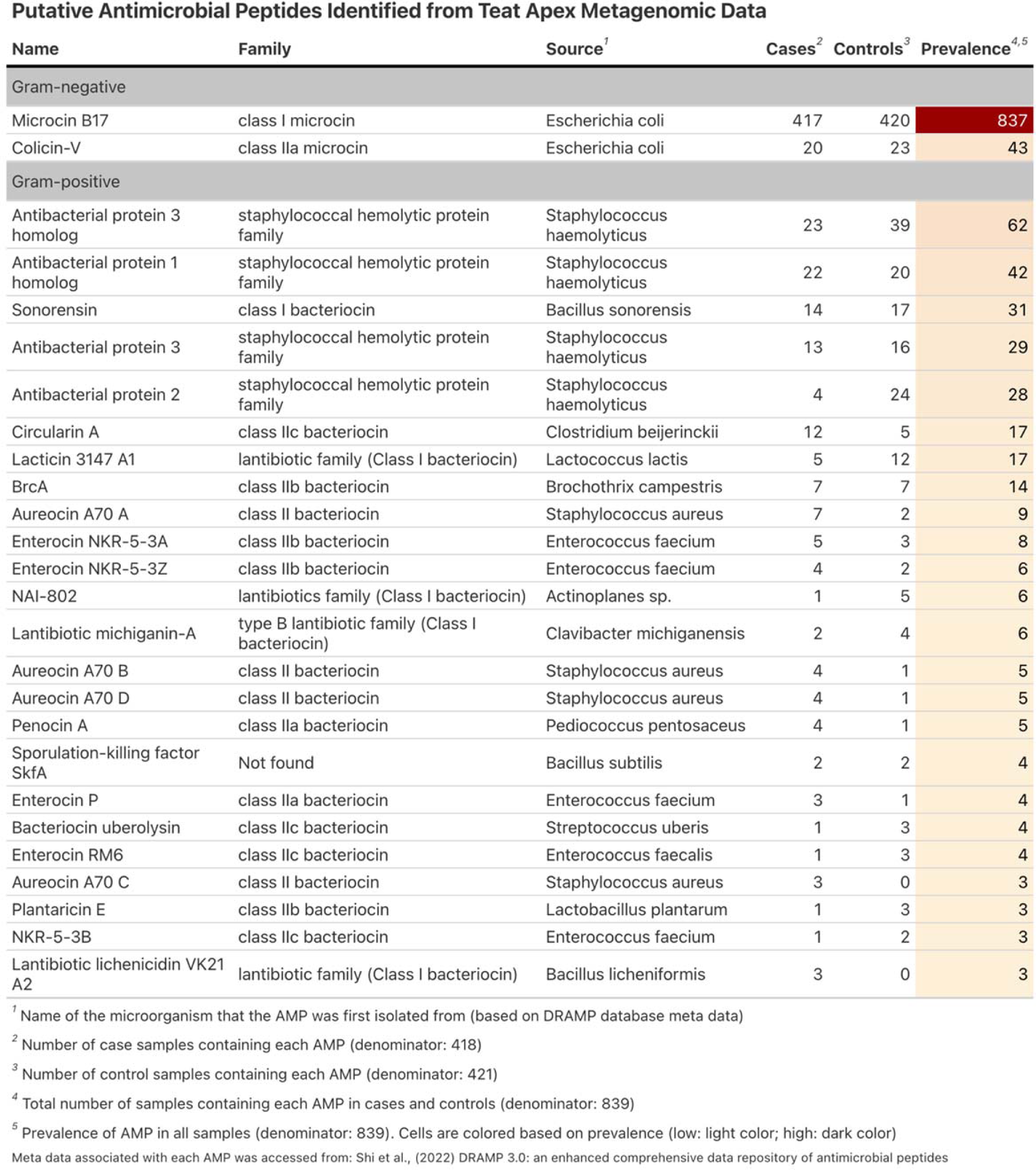
Antimicrobial peptides (AMPs) detected in the teat apex shotgun metagenomic data. AMPs were inferred by aligning translated sequence reads to the DRAMP protein database using PALADIN *(Westbrook et al., 2017)*. AMPs present in three or more samples are shown.

## DISCUSSION

We found that samples from cows testing positive for an *S. aureus* IMI after parturition had a significantly higher odds of *S. aureus* DNA within metagenomic data from the teat apex compared to cows that tested negative (OR = 38.95, 95% CI: 14.8-102.2). These findings are consistent with culture-based results reporting that nulliparous cows with teats culturing positive for *S. aureus* have a 3.34-times higher odds of testing positive for an *S. aureus* IMI after parturition compared to heifers with teats that were culture negative for *S. aureus* (Roberson et al., 1994). The same study also showed that the presence of *S. aureus* cultured from lacteal secretions collected prior to parturition was associated with 19.56-higher odds of having an *S. aureus* IMI after parturition (Roberson et al., 1994). A recent cross-sectional study of 287 multiparous cows in early (<90 DIM), mid (90-199 DIM), and late (≥200 DIM) lactation found that when the teat skin from a quarter was culture positive for *S. aureus*, the odds of the same quarter being culture positive for an *S. aureus* IMI was 7.8-times higher compared to quarters with teats that were not culture positive for *S. aureus* (Svennesen et al., 2019). A smaller cross-sectional study of 57 multiparous cows also found that teat skin samples that were culture positive for *S. aureus* had 4.5-times higher risk of *S. aureus* IMI in the same quarter compared to teat skin samples that were not culture positive for *S. aureus* (da Costa et al., 2014). However, the cross-sectional nature of the latter two study designs and the time periods at which they occurred (i.e., during lactation) precluded establishing temporality between *S. aureus* colonization of the teat skin and *S. aureus* IMIs. A strength of our study is that we attempted to address these issues by sampling the teat-skin up to 8 weeks prior to parturition and on weekly intervals thereafter. However, our ability to definitively establish temporal relationships between the teat apex microbiome and *S. aureus* IMIs was complicated by the fact that many of the observed post-calving IMIs occurred during the first week post-calving. Indeed, the median number of days after calving for when a case was identified was six, which would certainly afford microorganisms enough time to establish an *S. aureus* IMI. However, the high number of case samples with the presence of *S. aureus* DNA on the teat apex compared to controls suggests that IMIs could have been acquired prior to calving (**Fig. 3D**), as was observed by (Roberson et al., 1994). If this is true, then the route of exposure may have been due to “open quarters”, which (Krömker and Friedrich, 2009) observed in 60% of teat canals as far back as 60 days prior to parturition in a study of 84 Holstein heifers. Indeed, prepartum IMIs with both minor and major pathogens such as *S. aureus* were common among open quarters, as was the proportion of postpartum IMIs (77%) caused by pathogens observed during the prepartum period (Krömker and Friedrich, 2009). We also observed that animals with an *S. aureus* IMI had higher odds of having herpes virus DNA on the teat apex, specifically a herpes virus belonging to the *Herpesvirales* order (OTU1326) (OR = 2.09, 95% CI: 1.46-2.98) compared to animals without an *S. aureus* IMI. This is an interesting finding because herpes-associated viruses are thought to trigger formation of teat lesions that can act as a reservoir for bacterial colonization of the teat skin, thus leading to increased risk of acquiring an IMI (Wellenberg et al., 2002).

Previous culture-based studies of the teat apex have also reported protective effects of specific bacterial groups against major mastitis pathogens such as *S. aureus* (Woodward et al., 1987; De Vliegher et al., 2004). One popular bacterial group with the purported effect has been NAS. Consistent with previous studies of the teat apex, NAS DNA was highly prevalent in our study, with *S. chromogenes*, *S. devriesei*, *S. haemolyticus* and *S. auricularis* all attaining at least 45% prevalence across our sample set (Braem et al., 2013; De Visscher et al., 2016; Mahmmod et al., 2018). However, in contrast to observations reported in previous field and *in vitro* studies, we were not able to provide strong support for the hypothesis that the presence of NAS DNA on the teat apex confers protection against *S. aureus* colonization of the mammary gland (De Vliegher et al., 2004; Braem et al., 2014). In fact, there was no evidence of an association between the presence of DNA from *S. chromogenes* (OR = 0.97, 95% CI: 0.56-1.69), *S. devriesei* (OR = 0.78, 95% CI: 0.44-1.38) or *S. haemolyticus* (OR = 0.81, 95% CI: 0.47-1.38) on the teat apex and *S. aureus* IMI risk. Nor was there a difference in DNA abundance of these bacteria between cases and controls. In many cases, the inhibition of *S. aureus* by *S. chromogenes* in previous studies was observed within *in vitro* models (De Vliegher et al., 2004), which is not likely to be reflective of the teat apex environment. Indeed, recent concerns about the utility and reproducibility of *in vitro* experiments using culture models that are not reflective of the complex microbial environment under study has been raised within the life science community (Hirsch and Schildknecht, 2019). A meta-analysis investigating the association between the presence of minor pathogens (NAS and *C. bovis*) on major pathogen (*S. aureus*, *S. agalactiae*, *S. uberis*, *S. dysgalactiae*, *E. coli*) IMI risk found that, on average, there was no association in observational studies, but there was a significantly lower risk, on average, observed in experimental studies (Reyher et al., 2012). As the authors point out, heterogeneity between both observational and experimental studies do exist, which may explain the null findings observed in our study (Reyher et al., 2012). Many of the protective associations conferred by NAS have also been explored within milk, whereas our study focused on the skin, which may also explain the null associations observed for most NAS species investigated in our study. Finally, our analysis utilized metagenomic data, which can contain DNA from both live and dead organisms; whereas previous studies reporting on the protective effect of NAS utilized culture-based assays wherein only viable NAS bacteria were included.

Our study also identified several microorganisms on the teat apex that were associated with lower odds of acquiring an *S. aureus* IMI. The microorganisms with the largest protective effects included *Corynebacterium camporealensis*, *Corynebacterium maris*, *Kocuria polaris*, *Kocuria rosea*, *Dietzia alimantera*, *Cutibacterium acnes*, *Psychrobacter immobilis* and *Microbacterium phage min 1*. Many of these genera and species have been reported in the relatively few existing studies that utilize high-throughput sequencing to interrogate the teat apex microbiome of dairy cows (Braem et al., 2012; Verdier-Metz et al., 2012; Dean et al., 2021). However, little is known about the specific effects these microorganisms have on udder health, as most of these microorganisms have not been widely studied using traditional culture-based methods. The exception to this is *Cutibacterium acnes* (formerly *Propionibacterium acnes* and *Corynebacterium parvum* (Dinsmore et al., 1995)), which had a short history of being evaluated as an intramammary immunostimulant during the dry period to prevent new IMIs after parturition (Hogan et al., 1994) and as a treatment for chronic *S. aureus* IMIs (Dinsmore et al., 1995). The main takeaway from both studies was that *Cutibacterium acnes* cultured from milk samples had little to no effect on udder health outcomes (Hogan et al., 1994; Dinsmore et al., 1995). The positive effect observed in our study may be explained by the fact that *C. acnes* was found on the teat apex, and therefore may act differently in this environmental niche compared to inside the mammary gland itself. Bacteria belonging to the *Corynebacterium* genera are generally considered to be minor IMI pathogens, but their presence within the mammary gland has also been associated with a lower odds of *S. aureus* IMI in experimental studies, though findings are mixed (Reyher et al., 2012). Information about the specific genera within the *Corynebacterium* genus that we identified as being protective (i.e., *C. camporealensis* and *C. maris*) is scarce, as these bacteria have not been reported in most culture-based studies involving udder health. This is likely because many bacteria are not culturable, and the MALDI-TOF MS instrumentation used to assign taxonomic labels is limited to the taxa that are well-represented in reference databases. The microorganism with the strongest protective effect was *Microbacterium phage Min 1* (OR = 0.37, 95% CI: 0.25-0.53), a bacteriophage that has been found on plasmids within bacterial cells of the *Microbacterium* genus (Akimkina et al., 2007).

Indeed, 8 species of bacteria belonging to the *Microbacterium* genus were observed on the teat apex (**Supplementary Table 2**). Bacteria belonging to the *Microbacterium* genus have also been reported as common laboratory contaminants (Salter et al., 2014), but this seems to be an unlikely source in our study as we would expect an equal probability of these bacteriophage to be found in both cases and controls since our samples were processed using the same lot of laboratory reagents (adjusted risk in cases = 63.0% in cases, adjusted risk in controls 82.0%).

An unexpected finding of our metagenomic study was the difference in abundance of reads that aligned to the *Bos taurus* genome between cases and controls (**Fig. 1 G-J**). Because we sequenced DNA, it is impossible to determine what types of host cell populations were sequenced, but somatic cells such as keratinocytes (i.e., epidermal cells) are likely as we sampled skin (Nickoloff and Turka, 1993). Indeed, keratinocytes are the most common cell type found on skin, and they play a fundamental role in maintaining the integrity of the skin in the presence of microorganisms (Sørensen et al., 2003; Wang and Li, 2020; Mestrallet et al., 2021). Another type of somatic cell that may have been sampled are neutrophils, which are common within the mammary gland of dairy cows (Alhussien and Dang, 2018), especially during an intramammary infection (Rainard et al., 2018). In fact, the concentration of neutrophils and other somatic cells in bovine milk (termed “somatic cell count”) is a widely used indicator for udder health and mastitis screening (Reneau, 1986). Although we did not sample milk, neutrophils can also be found in skin when responding to *Staphylococcus aureus* infections in humans (Miller and Cho, 2011), which may explain the elevated abundance of *B. taurus* DNA in skin samples from cases in our study (**Fig. 1 I-J**). However, this may only provide a partial explanation, as *Bos taurus* DNA was consistently higher in samples from cases compared to controls throughout the entire sampling period. It may be tempting to attribute this finding to a technical artifact of our study design, but we found no evidence for this, as the elevated levels of *B. taurus* DNA in cases versus controls was found consistently across all sequencing batches (**Fig. 1G)** and there was no difference in raw sequencing depth between samples collected from cases and controls (**Fig. 1 B-C**). Although we observed a statistically significant batch effect, the effect size was small and unlikely to be a major contributing factor. Taken together, this finding appears to reflect a real biological phenomenon that deserves further attention.

In addition to exploring associations between the teat apex metagenome and *S. aureus* IMI risk, we also investigated the functional capacity of the teat apex microbiome to produce AMPs that could influence *S. aureus* IMI risk. Our metagenomic analysis identified 74 unique putative AMPs with a presumed bacterial origin based on annotation information provided in the DRAMP reference database (Shi et al., 2022). The most prevalent of these AMPs were Class I and II microcins, thought to be primarily produced by bacteria belonging to the *Enterobacteriaceae* family (Parker and Davies, 2022). Although we could not verify the specific bacterial origin of these AMPs, our finding that Microcin abundance was greater in controls compared to cases (*p* = 0.0014) is interesting, though their antibacterial effect is thought to be restricted to closely related bacteria, i.e., those within the *Enterobacteriaceae* family (Duquesne et al., 2007), of which *S. aureus* is not a member. We also identified putative AMPs from the staphylococcal hemolytic protein family, which are thought to originate from *Staphylococcus haemolyticus* (Watson et al., 1988; Takeuchi et al., 2005). Intriguingly, these AMPs were more prevalent in controls versus cases (crude prevalence of 7.8% versus 5.0%), suggesting a potential protective effect against *S. aureus* IMIs. *S. haemolyticus* belongs to the NAS group of bacteria that are thought to play a protective role in *S. aureus* IMI risk through the production of these types of molecules (Braem et al., 2014). It is important to note that our data suggest that bacteria from the teat apex have the functional capacity to produce these peptides, but as we identified them using an *in silico* and DNA-based approach, we must be cautious about interpreting these results – they are simply hypotheses. Currently, little is known about the role of these naturally occurring peptides on IMI risk, but further research on their effect using proteomic assays could be explored, especially considering the current need for alternative antimicrobial therapies within agricultural settings (Diez-Gonzalez, 2007).

Although we identified numerous microorganisms that were significantly associated with the presence of post-calving *S. aureus* IMIs, we also found that these results need to be interpreted within the context of their variance components, i.e., cows and farms. As an example, our model investigating the association between *S. aureus* DNA on the teat apex and postpartum *S. aureus* IMI risk found that there was more variation between cows (ICC: 0.32) than between farms (ICC: 0.10). However, this was not the case for a similar model investigating the association between *Pychrobacter immobilis* DNA and *S. aureus* IMI risk, in which the variation between cows (ICC: 0.04) was less than that between farms (ICC: 0.77). Indeed, the latter microbe was primarily isolated within a single farm (high farm ICC), while the former was isolated consistently across each farm (low farm ICC). This is visually apparent when comparing the abundances of these two specific microorganisms between cases and controls across each farm (**Fig. 4B**). More broadly, this implies that the teat apex microbiome is not likely to be homogenous in animals between farms, and that future studies of the teat apex microbiome should consider accounting for this biological effect when interpreting their results, as results with large effect sizes or significant associations cannot simply be generalized across all herds.

This certainly poses an interesting dilemma for future studies of the teat apex microbiome, as the investigator will need to think carefully about the number of herds to enroll and if enrolling multiple herds in similar locations or with similar management practices will mitigate the differences observed across farms.

## CONCLUSIONS

*Staphylococcus aureus* DNA on the teat apex was the strongest risk factor for *S. aureus* IMI after parturition and exhibited the largest differential abundance between cases and controls. Further research into the role of bacteriophage and other bacteria associated with lower *S. aureus* IMI risk could be explored to further understand their putative impact on udder health. Based on our results, the genera *Psychrobacter*, *Corynebacterium, Dietzia*, *Kocuria* and *Cutibacterium* and phages associated with *Microbacterium* may be promising candidates for this future research.

The continued exploration of AMPs (particularly Microcin B17 and AMPs produced by *S. haemolyticus*) could yield non-antibiotic alternatives for improved udder health.

## DATA AVAILABILITY

The raw sequence data generated in this study has been submitted to the Sequence Read Archive on NCBI (Accession: PRJNA984925).

## AUTHORS’ CONTRIBUTIONS

CD wrote initial versions of the manuscript, conducted formal analysis, generated data visualizations, and was responsible for software development. FPM, SMG and NRN provided valuable inputs on statistical analysis and bioinformatics. CD, FPM, TW, KS, AA, ED, LF, VFC, CB, BH, PP, VSM, LSC and NN were responsible for sample collection. BH, PP, VSM, LSC and NN supervised sample collection and farm enrollment. SMG provided mentoring and intellectual support to the lead author on this work. TR led and supervised all laboratory work conducted by CD, FPM, TW, VFC, and CB. NRN and LSC designed and acquired funding for this study. All authors provided input on the manuscript.

## ACKNOWLEDGEMENTS

We would like to thank the University of Minnesota Genomics Core (UMGC) for library preparation and sequencing support; the Minnesota Supercomputing Institute (MSI) for providing data storage and computational resources; the University of Minnesota Laboratory of Udder Health for bacteriology support; the farm personnel and owners who allowed us to work with them and their animals; and student volunteers who assisted with sample collection.

## REFERENCES

1. Aitchison, J. 1982. The Statistical Analysis of Compositional Data. Journal of the Royal Statistical Society: Series B (Methodological) 44:139–160. doi:10.1111/j.2517-6161.1982.tb01195.x.

2. Akimkina, T., C. Venien-Bryan, and J. Hodgkin. 2007. Isolation, characterization and complete nucleotide sequence of a novel temperate bacteriophage Min1, isolated from the nematode pathogen Microbacterium nematophilum. Res Microbiol 158:582–590. doi:10.1016/j.resmic.2007.06.005.

3. Alhussien, M.N., and A.K. Dang. 2018. Milk somatic cells, factors influencing their release, future prospects, and practical utility in dairy animals: An overview. Vet World 11:562–577. doi:10.14202/vetworld.2018.562-577.

4. Anderson, M.J. 2001. A new method for non-parametric multivariate analysis of variance. Austral Ecology 26:32–46. doi:10.1111/j.1442-9993.2001.01070.pp.x.

5. Andrews, T., D.A. Neher, T.R. Weicht, and J.W. Barlow. 2019. Mammary microbiome of lactating organic dairy cows varies by time, tissue site, and infection status. PLOS ONE 14:e0225001. doi:10.1371/journal.pone.0225001.

6. Barkema, H.W., M.J. Green, A.J. Bradley, and R.N. Zadoks. 2009. The role of contagious disease in udder health. J Dairy Sci 92:4717–4729. doi:10.3168/jds.2009-2347.

7. Barkema, H.W., Y.H. Schukken, and R.N. Zadoks. 2006. Invited Review: The Role of Cow, Pathogen, and Treatment Regimen in the Therapeutic Success of Bovine Staphylococcus aureus Mastitis. Journal of Dairy Science 89:1877–1895. doi:10.3168/jds.S0022-0302(06)72256-1.

8. Bates, D., M. Mächler, B. Bolker, and S. Walker. 2015. Fitting Linear Mixed-Effects Models Using lme4. Journal of Statistical Software 67:1–48. doi:10.18637/jss.v067.i01.

9. Beghini, F., L.J. McIver, A. Blanco-Míguez, L. Dubois, F. Asnicar, S. Maharjan, A. Mailyan, P. Manghi, M. Scholz, A.M. Thomas, M. Valles-Colomer, G. Weingart, Y. Zhang, M. Zolfo, C. Huttenhower, E.A. Franzosa, and N. Segata. 2021. Integrating taxonomic, functional, and strain- level profiling of diverse microbial communities with bioBakery 3. eLife 10:e65088. doi:10.7554/eLife.65088.

10. Benjamini, Y., and Y. Hochberg. 1995. Controlling the False Discovery Rate: A Practical and Powerful Approach to Multiple Testing. Journal of the Royal Statistical Society: Series B (Methodological) 57:289–300. doi:10.1111/j.2517-6161.1995.tb02031.x.

11. Braem, G., S. De Vliegher, B. Verbist, V. Piessens, E. Van Coillie, L. De Vuyst, and F. Leroy. 2013.Unraveling the microbiota of teat apices of clinically healthy lactating dairy cows, with special emphasis on coagulase-negative staphylococci.. J Dairy Sci 96:1499–1510. doi:10.3168/jds.2012-5493.

12. Braem, G., B. Stijlemans, W. Van Haken, S. De Vliegher, L. De Vuyst, and F. Leroy. 2014. Antibacterial activities of coagulase-negative staphylococci from bovine teat apex skin and their inhibitory effect on mastitis-related pathogens. J Appl Microbiol 116:1084–1093. doi:10.1111/jam.12447.

13. Braem, G., S. de Vliegher, B. Verbist, M. Heyndrickx, F. Leroy, and L. de Vuyst. 2012. Culture- independent exploration of the teat apex microbiota of dairy cows reveals a wide bacterial species diversity. Veterinary Microbiology 383–390. doi:10.1016/j.vetmic.2011.12.031.

14. Brooks, M., E., K. Kristensen, K. Benthem J.,van, A. Magnusson, C. Berg W., A. Nielsen, H. Skaug J., M. Mächler, and B. Bolker M. 2017. glmmTMB Balances Speed and Flexibility Among Packages for Zero-inflated Generalized Linear Mixed Modeling. The R Journal 9:378. doi:10.32614/RJ-2017-066.

15. da Costa, L.B., P.J. Rajala-Schultz, A. Hoet, K.S. Seo, K. Fogt, and B.S. Moon. 2014. Genetic relatedness and virulence factors of bovine Staphylococcus aureus isolated from teat skin and milk. Journal of Dairy Science 97:6907–6916. doi:10.3168/jds.2014-7972.

16. Cotter, P.D., C. Hill, and R.P. Ross. 2005. Bacteriocins: developing innate immunity for food. Nat Rev Microbiol 3:777–788. doi:10.1038/nrmicro1273.

17. De Visscher, A., S. Piepers, F. Haesebrouck, and S. De Vliegher. 2016. Teat apex colonization with coagulase-negative Staphylococcus species before parturition: Distribution and species-specific risk factors. J Dairy Sci 99:1427–1439. doi:10.3168/jds.2015-10326.

18. De Vliegher, S. 2003. Prepartum teat apex colonization with Staphylococcus chromogenes in dairy heifers is associated with low somatic cell count in early lactation. Veterinary Microbiology 92:245–252. doi:10.1016/S0378-1135(02)00363-2.

19. De Vliegher, S., L.K. Fox, S. Piepers, S. McDougall, and H.W. Barkema. 2012. Invited review: Mastitis in dairy heifers: nature of the disease, potential impact, prevention, and control. J Dairy Sci 95:1025–1040. doi:10.3168/jds.2010-4074.

20. De Vliegher, S., G. Opsomer, A. Vanrolleghem, L.A. Devriese, O.C. Sampimon, J. Sol, H.W. Barkema, F. Haesebrouck, and A. de Kruif. 2004. In vitro growth inhibition of major mastitis pathogens by Staphylococcus chromogenes originating from teat apices of dairy heifers. Veterinary Microbiology 101:215–221. doi:10.1016/j.vetmic.2004.03.020.

21. Dean, C.J., F. Peña-Mosca, T. Ray, B.J. Heins, V.S. Machado, P.J. Pinedo, L.S. Caixeta, and N.R. Noyes. 2022. Evaluation of Contamination in Milk Samples Pooled From Independently Collected Quarters Within a Laboratory Setting. Frontiers in Veterinary Science 9.

22. Dean, C.J., I.B. Slizovskiy, K.K. Crone, A.X. Pfennig, B.J. Heins, L.S. Caixeta, and N.R. Noyes. 2021. Investigating the cow skin and teat canal microbiomes of the bovine udder using different sampling and sequencing approaches. Journal of Dairy Science 104:644–661. doi:10.3168/jds.2020-18277.

23. Di Tommaso, P., M. Chatzou, E.W. Floden, P.P. Barja, E. Palumbo, and C. Notredame. 2017. Nextflow enables reproducible computational workflows. Nat Biotechnol 35:316–319. doi:10.1038/nbt.3820.

24. Diez-Gonzalez, F. 2007. Applications of Bacteriocins in Livestock.

25. Dinsmore, R.P., M.B. Cattell, R.D. Stevens, C.S. Gabel, M.D. Salman, and J.K. Collins. 1995. Efficacy of a Propionibacterium acnes Immunostimulant for Treatment of Chronic Staphylococcus aureus Mastitis. Journal of Dairy Science 78:1932–1936. doi:10.3168/jds.S0022-0302(95)76818-7.

26. Dohoo, I., W. Martin, and H. Stryhn. 2009. Veterinary Epidemiologic Research – Third Printing of the Second Edition | Ian Dohoo | Wayne Martin | Henrik Stryhn.

27. Duquesne, S., D. Destoumieux-Garzón, J. Peduzzi, and S. Rebuffat. 2007. Microcins, gene-encoded antibacterial peptides from enterobacteria. Nat Prod Rep 24:708–734. doi:10.1039/b516237h.

28. Ewels, P., M. Magnusson, S. Lundin, and M. Käller. 2016. MultiQC: summarize analysis results for multiple tools and samples in a single report. Bioinformatics 32:3047–3048. doi:10.1093/bioinformatics/btw354.

29. Falentin, H., L. Rault, A. Nicolas, D.S. Bouchard, J. Lassalas, P. Lamberton, J.-M. Aubry, P.-G. Marnet, Y. Le Loir, and S. Even. 2016. Bovine Teat Microbiome Analysis Revealed Reduced Alpha Diversity and Significant Changes in Taxonomic Profiles in Quarters with a History of Mastitis. Front Microbiol 7. doi:10.3389/fmicb.2016.00480.

30. Fang, R., B. Wagner, J.K. Harris, and S.A. Fillon. 2014. Application of zero-inflated negative binomial mixed model to human microbiota sequence data. PeerJ Inc.

31. Fernandes, A.D., J.N. Reid, J.M. Macklaim, T.A. McMurrough, D.R. Edgell, and G.B. Gloor. 2014. Unifying the analysis of high-throughput sequencing datasets: characterizing RNA-seq, 16S rRNA gene sequencing and selective growth experiments by compositional data analysis. Microbiome 2:15. doi:10.1186/2049-2618-2-15.

32. Ferronatto, J.A., F.N. Souza, A.M.M.P.D. Libera, S.D. Vliegher, A.D. Visscher, S. Piepers, M.G. Blagitz, and M.B. Heinemann. 2019. Inhibition of the growth of major mastitis-causing pathogens by non- aureus Staphylococcus isolates using the cross-streaking method. Arq. Bras. Med. Vet. Zootec. 71:1745–1749. doi:10.1590/1678-4162-11006.

33. Hirsch, C., and S. Schildknecht. 2019. In Vitro Research Reproducibility: Keeping Up High Standards. Frontiers in Pharmacology 10.

34. Hogan, J.S., K.L. Smith, D.A. Todhunter, P.S. Schoenberger, R.P. Dinsmore, M.B. Canttell, and C.S. Gabel. 1994. Efficacy of dry cow therapy and a Propionibacterium acnes product in herds with low somatic cell count. J Dairy Sci 77:3331–3337. doi:10.3168/jds.S0022-0302(94)77274-X.

35. Jahan, N.A., S.M. Godden, E. Royster, T.C. Schoenfuss, C. Gebhart, J. Timmerman, and R.C. Fink. 2021. Evaluation of the matrix-assisted laser desorption ionization time of flight mass spectrometry (MALDI-TOF MS) system in the detection of mastitis pathogens from bovine milk samples. Journal of Microbiological Methods 182:106168. doi:10.1016/j.mimet.2021.106168.

36. Kabera, F., J.-P. Roy, M. Afifi, S. Godden, H. Stryhn, J. Sanchez, and S. Dufour. 2021. Comparing Blanket vs. Selective Dry Cow Treatment Approaches for Elimination and Prevention of Intramammary Infections During the Dry Period: A Systematic Review and Meta-Analysis. Front Vet Sci 8:688450. doi:10.3389/fvets.2021.688450.

37. Krömker, V., and J. Friedrich. 2009. Teat canal closure in non-lactating heifers and its association with udder health in the consecutive lactation. Veterinary Microbiology 134:100–105. doi:10.1016/j.vetmic.2008.09.002.

38. Krueger, F., F. James, P. Ewels, E. Afyounian, and B. Schuster-Boeckler. 2021. FelixKrueger/TrimGalore: v0.6.7 - DOI via Zenodo. doi:10.5281/zenodo.5127899.

39. Lenth, R.V., P. Buerkner, M. Herve, M. Jung, J. Love, F. Miguez, H. Riebl, and H. Singmann. 2022. emmeans: Estimated Marginal Means, aka Least-Squares Means.

40. Li, H. 2013. Aligning sequence reads, clone sequences and assembly contigs with BWA-MEM. arXiv:1303.3997 [q-bio].

41. Li, H., B. Handsaker, A. Wysoker, T. Fennell, J. Ruan, N. Homer, G. Marth, G. Abecasis, and R. Durbin. 2009. The Sequence Alignment/Map format and SAMtools. Bioinformatics 25:2078–2079. doi:10.1093/bioinformatics/btp352.

42. Love, M.I., W. Huber, and S. Anders. 2014. Moderated estimation of fold change and dispersion for RNA-seq data with DESeq2. Genome Biology 15:550. doi:10.1186/s13059-014-0550-8.

43. Lüdecke, D., M.S. Ben-Shachar, I. Patil, P. Waggoner, and D. Makowski. 2021. performance: An R Package for Assessment, Comparison and Testing of Statistical Models. Journal of Open Source Software 6:3139. doi:10.21105/joss.03139.

44. Mahmmod, Y.S., I.C. Klaas, L. Svennesen, K. Pedersen, and H. Ingmer. 2018. Communications of Staphylococcus aureus and non-aureus Staphylococcus species from bovine intramammary infections and teat apex colonization. Journal of Dairy Science 101:7322–7333. doi:10.3168/jds.2017-14311.

45. Mallick, H., A. Rahnavard, L.J. McIver, S. Ma, Y. Zhang, L.H. Nguyen, T.L. Tickle, G. Weingart, B. Ren, E.H. Schwager, S. Chatterjee, K.N. Thompson, J.E. Wilkinson, A. Subramanian, Y. Lu, L. Waldron, J.N. Paulson, E.A. Franzosa, H.C. Bravo, and C. Huttenhower. 2021. Multivariable association discovery in population-scale meta-omics studies. PLOS Computational Biology 17:e1009442. doi:10.1371/journal.pcbi.1009442.

46. McCubbin, K.D., E. de Jong, T.J.G.M. Lam, D.F. Kelton, J.R. Middleton, S. McDougall, S.D. Vliegher, S. Godden, P.J. Rajala-Schultz, S. Rowe, D.C. Speksnijder, J.P. Kastelic, and H.W. Barkema. 2022. Invited review: Selective use of antimicrobials in dairy cattle at drying-off. Journal of Dairy Science 105:7161–7189. doi:10.3168/jds.2021-21455.

47. McMurdie, P.J., and S. Holmes. 2013. phyloseq: An R Package for Reproducible Interactive Analysis and Graphics of Microbiome Census Data. PLOS ONE 8:e61217. doi:10.1371/journal.pone.0061217.

48. Mestrallet, G., N. Rouas-Freiss, J. LeMaoult, N.O. Fortunel, and M.T. Martin. 2021. Skin Immunity and Tolerance: Focus on Epidermal Keratinocytes Expressing HLA-G. Frontiers in Immunology 12.

49. Miller, L.S., and J.S. Cho. 2011. Immunity against Staphylococcus aureus cutaneous infections. Nat Rev Immunol 11:505–518. doi:10.1038/nri3010.

50. Nickoloff, B.J., and L.A. Turka. 1993. Keratinocytes: key immunocytes of the integument.. Am J Pathol 143:325–331.

51. Oksanen, J., G.L. Simpson, F.G. Blanchet, R. Kindt, P. Legendre, P.R. Minchin, R.B. O’Hara, P. Solymos, M.H.H. Stevens, E. Szoecs, H. Wagner, M. Barbour, M. Bedward, B. Bolker, D. Borcard, G. Carvalho, M. Chirico, M.D. Caceres, S. Durand, H.B.A. Evangelista, R. FitzJohn, M. Friendly, B. Furneaux, G. Hannigan, M.O. Hill, L. Lahti, D. McGlinn, M.-H. Ouellette, E.R. Cunha, T. Smith, A. Stier, C.J.F.T. Braak, and J. Weedon. 2022. vegan: Community Ecology Package.

52. Parker, J.K., and B.W. Davies. 2022. Microcins reveal natural mechanisms of bacterial manipulation to inform therapeutic development. Microbiology 168:001175. doi:10.1099/mic.0.001175.

53. Paulson, J.N., O.C. Stine, H.C. Bravo, and M. Pop. 2013. Differential abundance analysis for microbial marker-gene surveys. Nat Methods 10:1200–1202. doi:10.1038/nmeth.2658.

54. Pol, M., and P.L. Ruegg. 2007a. Relationship between antimicrobial drug usage and antimicrobial susceptibility of gram-positive mastitis pathogens. J Dairy Sci 90:262–273. doi:10.3168/jds.S0022-0302(07)72627-9.

55. Pol, M., and P.L. Ruegg. 2007b. Treatment practices and quantification of antimicrobial drug usage in conventional and organic dairy farms in Wisconsin. J. Dairy Sci. 90:249–261. doi:10.3168/jds.S0022-0302(07)72626-7.

56. Rainard, P. 2017. Mammary microbiota of dairy ruminants: fact or fiction?. Veterinary Research 48:25. doi:10.1186/s13567-017-0429-2.

57. Rainard, P., G. Foucras, J.R. Fitzgerald, J.L. Watts, G. Koop, and J.R. Middleton. 2018. Knowledge gaps and research priorities in Staphylococcus aureus mastitis control. Transboundary and Emerging Diseases 65:149–165. doi:10.1111/tbed.12698.

58. Reneau, J.K. 1986. Effective Use of Dairy Herd Improvement Somatic Cell Counts in Mastitis Control. Journal of Dairy Science 69:1708–1720. doi:10.3168/jds.S0022-0302(86)80590-2.

59. Reyher, K.K., D. Haine, I.R. Dohoo, and C.W. Revie. 2012. Examining the effect of intramammary infections with minor mastitis pathogens on the acquisition of new intramammary infections with major mastitis pathogens—A systematic review and meta-analysis. Journal of Dairy Science 95:6483–6502. doi:10.3168/jds.2012-5594.

60. Ripley, B., B. Venables, D.M. Bates, K.H. (partial port ca 1998), A.G. (partial port ca 1998), and D. Firth. 2023. MASS: Support Functions and Datasets for Venables and Ripley’s MASS.

61. Roberson, J.R., L.K. Fox, D.D. Hancock, J.M. Gay, and T.E. Besser. 1994. Ecology of Staphylococcus aureus Isolated from Various Sites on Dairy Farms1. Journal of Dairy Science 77:3354–3364. doi:10.3168/jds.S0022-0302(94)77277-5.

62. Ruegg, P.L. 2009. Management of mastitis on organic and conventional dairy farms. J Anim Sci 87:43–55. doi:10.2527/jas.2008-1217.

63. Ruegg, P.L. 2017. A 100-Year Review: Mastitis detection, management, and prevention. Journal of Dairy Science 100:10381–10397. doi:10.3168/jds.2017-13023.

64. Salter, S.J., M.J. Cox, E.M. Turek, S.T. Calus, W.O. Cookson, M.F. Moffatt, P. Turner, J. Parkhill, N.J. Loman, and A.W. Walker. 2014. Reagent and laboratory contamination can critically impact sequence-based microbiome analyses. BMC Biology 12:87. doi:10.1186/s12915-014-0087-z.

65. Schreiner, D.A., and P.L. Ruegg. 2003. Relationship Between Udder and Leg Hygiene Scores and Subclinical Mastitis. Journal of Dairy Science 86:3460–3465. doi:10.3168/jds.S0022-0302(03)73950-2.

66. Shi, G., X. Kang, F. Dong, Y. Liu, N. Zhu, Y. Hu, H. Xu, X. Lao, and H. Zheng. 2022. DRAMP 3.0: an enhanced comprehensive data repository of antimicrobial peptides. Nucleic Acids Research 50:D488–D496. doi:10.1093/nar/gkab651.

67. Sordillo, L.M., K. Shafer-Weaver, and D. DeRosa. 1997. Immunobiology of the Mammary Gland. Journal of Dairy Science 80:1851–1865. doi:10.3168/jds.S0022-0302(97)76121-6.

68. Sørensen, O.E., J.B. Cowland, K. Theilgaard-Mönch, L. Liu, T. Ganz, and N. Borregaard. 2003. Wound Healing and Expression of Antimicrobial Peptides/Polypeptides in Human Keratinocytes, a Consequence of Common Growth Factors1. The Journal of Immunology 170:5583–5589. doi:10.4049/jimmunol.170.11.5583.

69. Svennesen, L., S.S. Nielsen, Y.S. Mahmmod, V. Krömker, K. Pedersen, and I.C. Klaas. 2019. Association between teat skin colonization and intramammary infection with Staphylococcus aureus and Streptococcus agalactiae in herds with automatic milking systems. Journal of Dairy Science 102:629–639. doi:10.3168/jds.2018-15330.

70. Takeuchi, F., S. Watanabe, T. Baba, H. Yuzawa, T. Ito, Y. Morimoto, M. Kuroda, L. Cui, M. Takahashi, A. Ankai, S. Baba, S. Fukui, J.C. Lee, and K. Hiramatsu. 2005. Whole-Genome Sequencing of Staphylococcus haemolyticus Uncovers the Extreme Plasticity of Its Genome and the Evolution of Human-Colonizing Staphylococcal Species. J Bacteriol 187:7292–7308. doi:10.1128/JB.187.21.7292-7308.2005.

71. Verdier-Metz, I., G. Gagne, S. Bornes, F. Monsallier, P. Veisseire, C. Delbès-Paus, and M.-C. Montel. 2012. Cow Teat Skin, a Potential Source of Diverse Microbial Populations for Cheese Production. Appl. Environ. Microbiol. 78:326–333. doi:10.1128/AEM.06229-11.

72. Wang, J.-N., and M. Li. 2020. The Immune Function of Keratinocytes in Anti-Pathogen Infection in the Skin. International Journal of Dermatology and Venereology 3:231. doi:10.1097/JD9.0000000000000094.

73. Watson, D.C., M. Yaguchi, J.G. Bisaillon, R. Beaudet, and R. Morosoli. 1988. The amino acid sequence of a gonococcal growth inhibitor from Staphylococcus haemolyticus. Biochem J 252:87–93. doi:10.1042/bj2520087.

74. Wellenberg, G.J., W.H.M. van der Poel, and J.T. Van Oirschot. 2002. Viral infections and bovine mastitis: a review. Vet Microbiol 88:27–45. doi:10.1016/s0378-1135(02)00098-6.

75. Westbrook, A., J. Ramsdell, T. Schuelke, L. Normington, R.D. Bergeron, W.K. Thomas, and M.D. MacManes. 2017. PALADIN: protein alignment for functional profiling whole metagenome shotgun data. Bioinformatics 33:1473–1478. doi:10.1093/bioinformatics/btx021.

76. Wickham, H. 2009. Ggplot2: Elegant Graphics for Data Analysis. 2nd ed. Springer Publishing Company, Incorporated.

77. Wood, D.E., J. Lu, and B. Langmead. 2019. Improved metagenomic analysis with Kraken 2. Genome Biology 20:257. doi:10.1186/s13059-019-1891-0.

78. Wood, D.E., and S.L. Salzberg. 2014. Kraken: ultrafast metagenomic sequence classification using exact alignments. Genome Biology 15:R46. doi:10.1186/gb-2014-15-3-r46.

79. Woodward, W.D., T.E. Besser, A.C. Ward, and L.B. Corbeil. 1987. In vitro growth inhibition of mastitis pathogens by bovine teat skin normal flora.. Can J Vet Res 51:27–31.

80. Zhang, X., Y.-F. Pei, L. Zhang, B. Guo, A.H. Pendegraft, W. Zhuang, and N. Yi. 2018. Negative Binomial Mixed Models for Analyzing Longitudinal Microbiome Data. Frontiers in Microbiology 9.

81. Zhang, X., and N. Yi. 2020. NBZIMM: negative binomial and zero-inflated mixed models, with application to microbiome/metagenomics data analysis. BMC Bioinformatics 21:488. doi:10.1186/s12859-020-03803-z.

